# Identifying potential keystone microbes from co-occurrence networks in the Gulf of Alaska

**DOI:** 10.1101/2025.06.12.659413

**Authors:** Megan Brauner, Jacob Cohen, Brandon R. Briggs, Gwenn M. M. Hennon

## Abstract

The Northern Gulf of Alaska (NGA) is a highly productive and diverse marine ecosystem. Differences in nutrient supply and physical circulation between nearshore and offshore waters in the NGA result in a mosaic of water masses with distinct biogeochemical signatures. We hypothesized that microbial communities in these regions not only differ in composition but also in the ecological interaction networks they support. We used amplicon sequencing of the 16S (V4) and 18S (V9) rRNA genes to characterize the microbial community differences between nearshore, continental shelf, and offshore regions in the NGA in summers 2018-2021. We observed significantly different community assemblages by region (MRPP, p = 0.001), with higher relative abundances and cell counts of heterotrophic bacteria and *Synechococcus* nearshore, elevated Alphaproteobacteria and SAR11 clades offshore, and greater dinoflagellates and Spirotrich ciliates on the shelf. Co-occurrence networks of operational taxonomic units (OTUs) of prokaryotes and eukaryotes were constructed for each region using statistically significant correlations (Spearman rank >0.8, Bonferroni corrected p < 0.05). Overall, the offshore network had higher centralization (0.331) and density (0.112), indicating higher connectivity and therefore more potential interactions compared to the shelf (0.191, 0.069) and nearshore (0.165, 0.041) networks. The nearshore network was characterized by higher proportions of potentially parasitic taxa such as *Cryothecomonas aestivalis*, Syndiniales Dino Group I, and MAST-1C and parasitoid bacteria Bdellovibrio and like organisms, suggesting that nearshore conditions may increase parasitoid/predator success through increased contact rates. Significant correlations between cryptophyte *Plagioselmis prolonga* and ciliate Oligotrichida were identified in all three regions, supporting previous findings that kleptoplasty is likely an important strategy across the NGA. Eukaryotic taxa that had the highest degree centrality across all three regions; *P. prolonga* and Phaeocystis are known to be mixotrophs, suggesting a role for bacterivory in forging a high number of interactions between protists and bacteria. Understanding these potential keystone microbes and their interactions is crucial for a greater understanding of ecosystem stability within the NGA.

## 1 Introduction

Oceans are experiencing rapid changes in temperature, stratification, and nutrient input because of climate change (Behrenfeld et al., 2006, 2006; Boyd et al., 2008; Hutchins and Fu, 2017). High-latitude regions, particularly the Arctic, are warming at a rate of approximately two to three times faster than the global average (Serreze et al., 2009; Screen and Simmonds, 2010; Cohen et al., 2014). Marine microbes make up more than 90% of ocean biomass and play critical roles in marine food webs, primary and secondary production, and the flow of energy and nutrients, making them significant drivers of biogeochemical cycles and environmental processes (Abirami et al., 2021). However, increased sea surface temperatures has shown to decrease microbial biodiversity (Hutchins and Fu, 2017; Abirami et al., 2021; Cohen, 2022) and have the potential to alter important microbial interactions and distributions (Cram et al., 2015; Fuhrman et al., 2015; Lima-Mendez et al., 2015; Worden et al., 2015).

The ocean’s large microbial populations and diversity likely support a complex network of interactions, from mutualism to antagonism, that shape marine ecosystems (Amin et al., 2012; Faust and Raes, 2012; Braga et al., 2016; Arandia-Gorostidi et al., 2022). Parasitoid bacteria like Bdellovibrio and like organisms (BALOs) can influence community structure by preying on small Gram-negative bacteria and helping regulate their populations (Sockett, 2009; Negus et al., 2017; Cohen et al., 2021; Mookherjee and Jurkevitch, 2022). Kleptoplasty, where microbes steal plastids from prey to temporarily acquire photosynthesis, is another important interaction, seen in marine ecosystems involving *Teleaulax amphioxeia*, *Mesodinium rubrum*, *Dinophysis*, and ciliates like *Strombidium* (Stoecker and Silver, 1990; McManus et al., 2012; Rial et al., 2012; Rusterholz et al., 2017; Cruz and Cartaxana, 2022). At the same time, mutualistic interactions are essential for maintaining microbial community structure and ecosystem services. For example, *Prochlorococcus* relies on helper bacteria like *Alteromonas* to remove reactive oxygen species (Morris et al., 2008, 2011; Hennon et al., 2018). These relationships are critical for ecosystem function, and shifts in ocean chemistry due to climate change could potentially destabilize these interactions (Hennon et al., 2018).

Although microbial interactions are important in shaping marine ecosystems, they are difficult to identify (Berry and Widder, 2014; Cagua et al., 2019; Matchado et al., 2021; Wang et al., 2021). Many marine microbes remain uncultured, and most of what is known about the diversity and distribution of these microbial communities comes from molecular techniques such as amplicon sequencing of the 16S and 18S rRNA genes (Hofer, 2018; Bodor et al., 2020). Network-based analytical approaches are useful for identifying potential microbial interactions based on co-occurrence or co-abundances of microbial taxa (Fuhrman, 2009; Needham et al., 2013; Berry and Widder, 2014; Cui et al., 2019; Li et al., 2021; Matchado et al., 2021; Costas-Selas et al., 2024). Using a co-occurrence network-based approach, significant correlations between microbial taxa are interpreted as potential interactions between those pairs (González et al., 2010; Berry and Widder, 2014; Cagua et al., 2019). Additionally, node-specific and global network measurements can provide statistical measurements about differences in putative interaction networks that help to uncover differences between co-occurrence networks’ structure and function (Faust and Raes, 2012; Berry and Widder, 2014; Cagua et al., 2019). These methods can also identify microbial taxa with high numbers of potential interactions within the system, termed keystone microbes, whose removal could disproportionately affect network stability (González et al., 2010; Cagua et al., 2019; Matchado et al., 2021).

In this study, we examined how microbial communities, co-occurrence networks and keystone microbes differ by region in the Northern Gulf of Alaska (NGA) Longterm Ecological Research (LTER) site. The NGA LTER represents an important subarctic region in the North Pacific Ocean that supports culturally and economically important fisheries (Holen, 2014; Szymkowiak, 2020). However, there are few published datasets examining microbial community diversity and dynamics, particularly for prokaryotes. Variations in water masses and high seasonality in primary production result in a mosaic of nutrient regions across the NGA. Nearshore communities are influenced by the Alaska Coastal Current (ACC), which is driven by along-shore winds and characterized by a low-salinity core from freshwater inputs (Strom et al., 2006; Stabeno et al., 2016). These nearshore waters are high in micronutrients such as iron, but are often limited by nitrate (Strom et al., 2006; Aguilar-Islas et al., 2016; Stabeno et al., 2016). The Alaska Current – Alaskan Stream is a cyclonic boundary current along the shelf break/slope that together with mesoscale eddies often found moving westward in this region (Ladd et al., 2007) promote mixing of offshore iron-deplete, but nitrate-replete waters with shelf waters, which in summer tend to be replete in iron relative to nitrate, creating patches that can support higher primary production (Aguilar-Islas et al., 2016; Stabeno et al., 2016; Coyle et al., 2019). Natural gradients of iron and macronutrients in the NGA create contrasting nutrient-limited conditions for microbial communities and their interactions. Based on these nutrient differences, we hypothesized that nearshore regions with their abundant micronutrients will exhibit lower network connectivity and more antagonistic interactions, as microbes can access micronutrients more readily without the need for complex relationships. In contrast, offshore regions, which are macronutrient-replete, but often iron-limited, will show higher network connectivity, as microbial communities in these areas will rely on more intricate and cooperative interactions to access essential micronutrients.

## 2 Methods

### 2.1 Sample Site

Water samples for DNA and flow cytometry analysis were collected within the NGA LTER site at various depths, including surface, ten meters, and at the depth of the deep chlorophyll maximum (DCM) (Cohen, 2022). Three transects: the Seward line, Middleton Island line, and Kodiak Island line were sampled to characterize the spatial variation of water properties throughout the NGA (Figure 1). Sampling of the Seward and Middleton Island lines was conducted in summers 2018-2021 while the Kodiak Island line was sampled summers 2018-2019 and 2021 (Supplemental Table 1). The 2018 cruise was conducted onboard *R/V Woldstad* and the other cruises were onboard *R/V Sikuliaq*. Regions were *a priori* selected based on distance from shore and their proximity to currents and bathymetric features (Figure 1). Nearshore stations were within 30 nautical miles or 55.6 km from shore in a region influenced by the ACC (Stabeno et al., 2016). Shelf stations greater than 30 nautical miles from shore and over the continental shelf as defined here as areas shallower than 300 meters depth. Offshore stations were defined as those over the continental slope (> 300 m bottom depth) and are influenced by the Alaska Current - Alaskan Stream and mesoscale eddies (Okkonen et al., 2001; Strom et al., 2010; Coyle et al., 2019). The map of sample locations was produced using Ocean Data View (Schlitzer, 2022).

**Figure 1.**
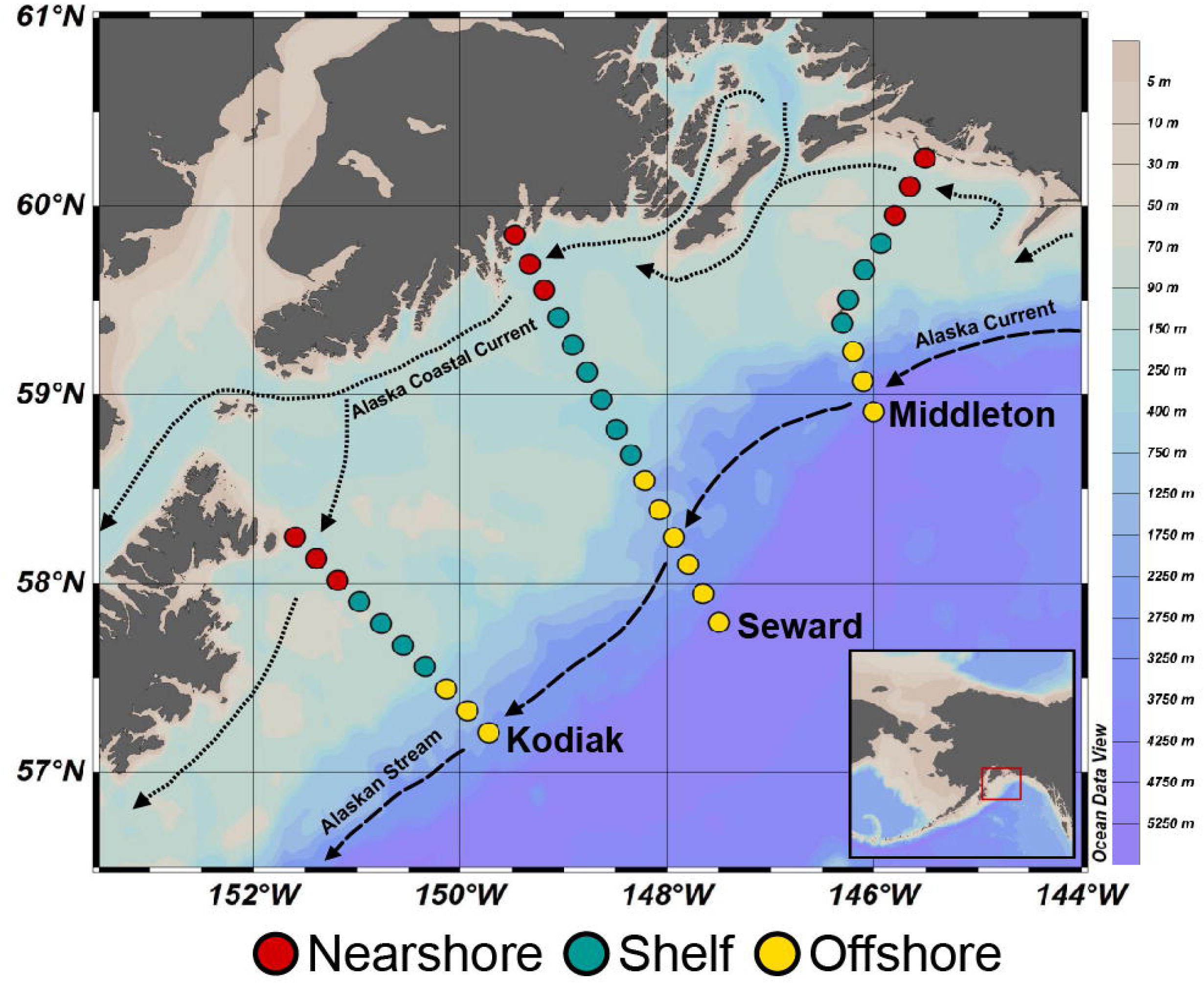
Stations for the Northern Gulf of Alaska (NGA). Sample categories based on location are differentiated by color; nearshore (red), shelf (teal), and offshore (yellow). Approximate locations of currents are indicated with dashed lines with directions indicated by arrows. Background color represents ocean bathymetry.

### 2.2 Flow Cytometry

Whole seawater was collected for flow cytometry in cryovials and preserved with a final concentration of 0.5% glutaraldehyde. Samples were fixed in the dark for ten minutes, flash-frozen in liquid nitrogen, and stored at −80°C until analysis. EasyCheck Beads were used to calibrate the flow cytometer, following a 20x dilution and running three replicates (Cytek, Fremont, USA). For DNA analysis, samples were stained with SYBR Green (100x dilution), incubated in the dark for ten minutes, and analyzed alongside unstained controls using a Guava 5ST easyCyte flow cytometer equipped with a 488-nm laser and detectors for side scatter, forward scatter, red, green, and yellow fluorescence (Luminex, Austin, USA). Following known optical properties, heterotrophic bacteria, and *Synechococcus* were manually gated using Guava InCyte (Luminex, Austin, USA).

### 2.3 Gene sequencing and bioinformatic analysis

Approximately 1-5 liters of seawater was filtered through a 0.2 µm pore Sterivex filter (Fisher Scientific, Hampton, USA) and frozen at −80°C until extraction. DNA was extracted from Sterivex filters using DNeasy PowerWater kit (Qiagen, Valencia, CA, USA) following the manufacturer’s protocol. Microbial community composition was analyzed by amplifying the 16S rRNA and 18S rRNA gene sequences using TaggiMatrix primer multiplexing (Glenn et al., 2019). Prokaryotic communities were amplified using the 16S (V4 region) 515F and 806R primers (Parada et al., 2016). Eukaryotic communities were amplified using the 18S (V9 region) 1389F and 1510R primers (Amaral-Zettler et al., 2009). Samples were sequenced using the Illumina MiSeq Platform (Illumina, San Diego, USA). A total of 232 prokaryotic and 198 eukaryotic samples were sequenced (PRJNA887083).

Raw sequences from Illumina were demultiplexed using Mr. Demuxy and processed using QIIME2 (version 2020.8) (Cock et al., 2009; Bolyen et al., 2019). Sequences were quality filtered (Phred score of 20) and denoised using DADA2 (Callahan et al., 2016). Operational taxonomic units (OTUs) were de novo clustered to 97% nucleotide identity and assigned taxonomy with the feature-classifier (Bokulich, 2018) using the Silva version 138 and PR^2^ reference databases (Guillou et al., 2012; Quast et al., 2013). Non-target 16S and 18S sequences, such as those identified as metazoan, chloroplast, and mitochondrial sequences were removed before further analysis. Samples were rarefied to a depth of 5,000 reads for the 16S rRNA gene and 2,500 reads for the 18S rRNA gene.

### 2.4 Co-occurrence Network Analysis

Microbial taxon-taxon co-occurrence networks were constructed for samples with significant Spearman rank correlations (ρ > 0.8) corrected for multiple testing (Bonferroni p < 0.05) in R using the Hmisc package (Harrell Jr and Dupont, 2019). Networks were visualized in Cytoscape (Shannon et al., 2003) and subnetworks and a merged network were constructed in Cytoscape using MetScape (Basu et al., 2017). Nodes represent all OTUs with significant correlations, which are represented by edges. Network and node-based measurements were calculated using the NetworkAnalyzer (Assenov et al., 2008). Degree centrality (connectivity) was calculated as the number of edges linked to each node (Diestel, 2005). Potential keystone microbes were identified as the top ten OTUs with the highest degree centrality values in each network. Closeness centrality of each node was calculated as the reciprocal of the average shortest path length (Newman, 2003), where the average shortest path length is the mean length of the shortest path between the node and all other nodes. Co-occurrence fragmentation (*f*) was calculated for each network as the ratio of disconnected subgraphs (CL) to the overall number of nodes (N) in each network (Berry and Widder, 2014; Yang et al., 2024). Where *f* can range from zero to one with zero being the most stable and 1 being the most unstable (Berry and Widder, 2014; Yang et al., 2024). A full set of network statistic definitions can be found in Supplemental Table 2.

### 2.5 Statistical Analysis

Non-metric multidimensional scaling ordination (NMDS) and significance of the multiple regression using environmental fit (envfit) with Bonferroni correction were conducted to test significant differences in microbial communities using the R package vegan (version 2.6-4, R version 4.2.1) (Dixon, 2003). Multiple response permutation procedure (MRPP) was conducted to test significant differences between microbial community composition using vegan (version 2.4-2) (Dixon, 2003). Differences in network measurements between sampling regions were compared using the non-parametric Wilcoxon rank sum test with the ggpubr package (Kassambara, 2020).

## 3 Results

### 3.1 Microbial community structure

Summer spatial variability in microbial communities and their interactions across the Northern Gulf of Alaska was investigated using seawater samples collected from three regions: nearshore, shelf, and offshore (Figure 1). DNA was extracted and amplified for the 16S and 18S rRNA genes for taxonomic identification of microbial communities. Amplicon sequencing of the 16S rRNA gene produced 5,080,643 sequences from 232 samples. After rarefying samples to a depth of 5,000 reads, 175 samples remained with a total of 875,000 sequences. 16S sequences were clustered into 1,863 operational taxonomic units (OTUs) which were assigned to 76 phyla. Amplicon sequencing of the 18S rRNA gene produced 3,484,572 sequences from 198 samples. After rarefying the samples to a depth of 2,500 reads, 145 samples remained with a total of 362,500 sequences. 18S sequences were clustered into 1165 OTUs which were assigned to 24 phyla.

Both prokaryotic (16S) and eukaryotic (18S) communities differed significantly by region (Figure 2, Bonferroni adjusted p < 0.05). The robustness of regional differences in community composition were further tested with multiple response permutation procedure (MRPP) and communities were found to differ significantly by region in both prokaryotes (p = 0.001) and eukaryotes (p = 0.001). Additionally, shifts in prokaryotic and eukaryotic communities were significantly correlated with environmental variables: temperature, salinity, nitrate, and silicic acid concentrations (Figure 2, Bonferroni adjusted, p < 0.05). MRPP analysis also showed significant differences with depth and year for both prokaryotes and eukaryotes. Previous research has confirmed significant interannual differences in microbial communities in the NGA (Cohen, 2022).

**Figure 2.**
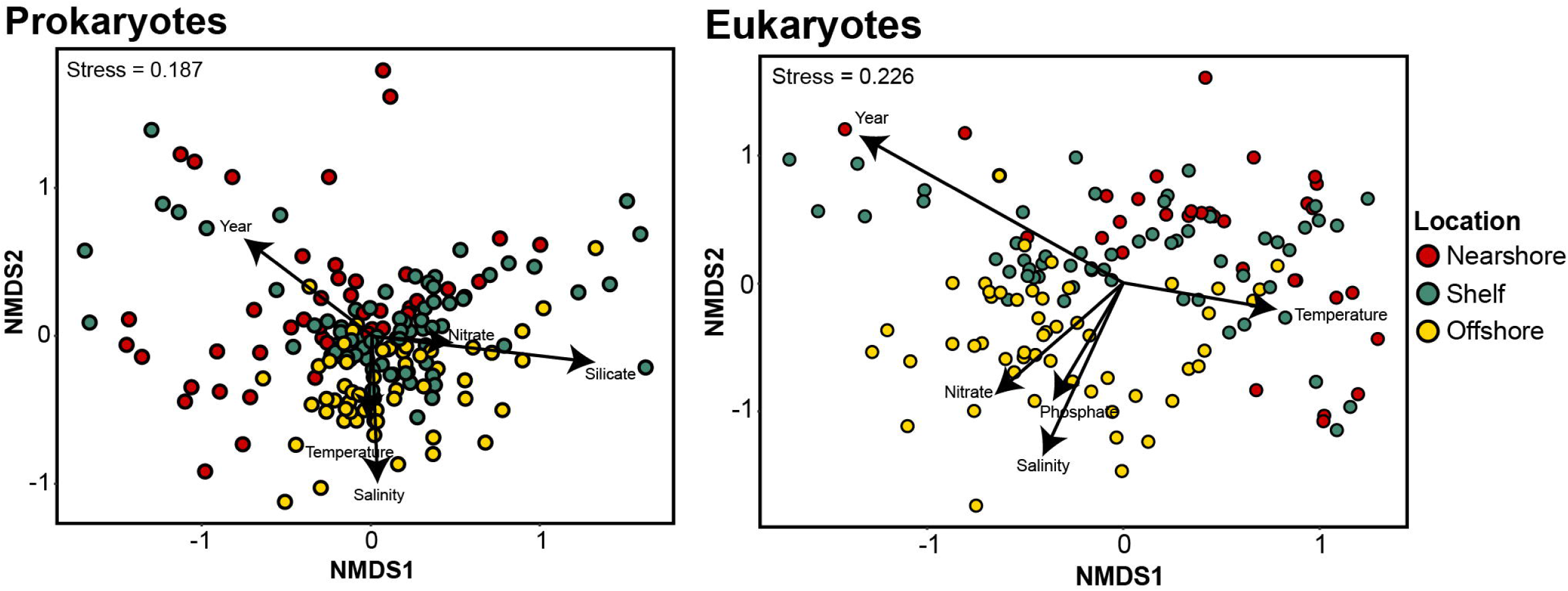
Nonmetric multidimensional scaling (NMDS) ordinations of variation in prokaryotic and eukaryotic water community structure. Color represents sample locations: Nearshore (red), Shelf (teal), and Offshore (yellow). Vectors represent significant correlations (p < 0.05) with environmental variables.

Shifts in the prokaryotic community were characterized by shifts in Alphaproteobacteria and Gammaproteobacteria. Alphaproteobacteria were the most abundant offshore (19.58%) and least abundant nearshore (14.45%) (Figure S1A). *Pelagibacter ubique* (SAR11), a prominent Alphaproteobacteria, varied by region, with clades I and II highest nearshore, clade IV highest offshore, and lowest levels on the shelf (Figure S1C). Gammaproteobacteria were most abundant nearshore (21.91%) and less abundant offshore and shelf. SAR86 within Gammaproteobacteria had the highest abundance offshore. Bdellovibrionota, also known as Bdellovibrio and like organisms (BALOs), remained stable between nearshore and offshore regions (2.53%).

Shifts in summertime eukaryotic communities were characterized by changes in dinoflagellates, diatoms, and prymnesiophytes. Dinoflagellates (phylum Dinoflagellata) were most abundant on the shelf (42.59%), with slightly lower relative abundance offshore (41.73%) and nearshore (37.07%) (Figure S1B). Within this group, the genus *Dinophysis*, a mixotrophic dinoflagellate, showed highest relative abundance offshore and lowest nearshore. The class Syndiniales, composed of parasitic dinoflagellates, was the most abundant across all regions. Group I Syndiniales were more common nearshore, while Groups II, III, and unclassified Syndiniales were more abundant offshore (Figure S1D). Gyrista, a recently defined phylum that includes diatoms and other stramenopiles, was most abundant offshore (24.57%), with lower relative abundance nearshore (13.59%) and on the shelf (12.30%). Within this group, pennate diatoms (class Bacillariophyceae) were more abundant nearshore (5.79%) and were primarily composed of the genus *Pseudo-nitzschia* (Figure S1E).

### 3.2 Differences in network statistics by region

Differences in microbial interactions by region were investigated by constructing microbial co-occurrence networks for each region using OTUs as nodes connected by statistically significant Spearman rank correlations (edges) (Figure 3; Table 1). The nearshore network contained the greatest number of nodes (n = 352), followed closely by the shelf network (n = 350), while the offshore network had the fewest (n = 226). In contrast, the shelf network had the highest number of edges (n = 1682), with the most positive correlations (n = 1552). The nearshore network had slightly fewer total edges (n = 1288), but still a high number of positive correlations (n = 1139). The offshore network had the fewest edges overall (n = 1127) and the highest proportion of negative correlations (n = 242). Eukaryotes dominated all three networks, representing the largest proportion of nodes in each region (Figure 3).

**Figure 3.**
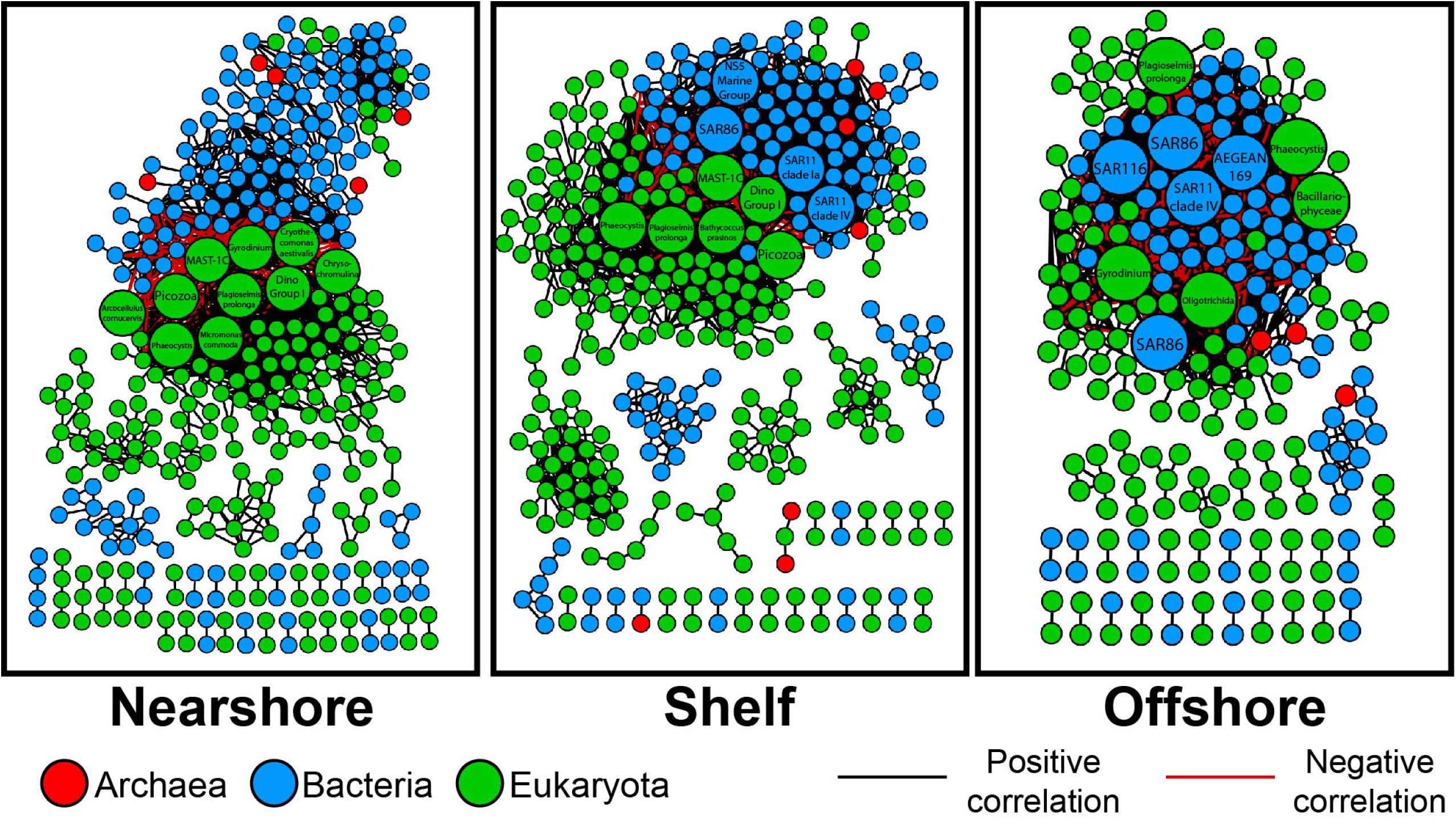
Co-occurrence networks of operational taxonomic units (OTUs) (nodes) with statistically significant Spearman rank correlations (> 0.8, Bonferroni corrected p < 0.05) including positive (black) and negative (red) (edges). Nodes are color-coded by taxonomy; Archaea (red), Bacteria (blue), and Eukaryota (green). Larger nodes have the highest degree centrality.

**Table 1.** Co-occurrence network statistics for Nearshore, Shelf, and Offshore networks.

| Network Statistic | Nearshore | Shelf | Offshore |
| --- | --- | --- | --- |
| Nodes | 352 | 350 | 226 |
| Edges | 1288 | 1682 | 1127 |
| Positive Edges | 1139 | 1552 | 885 |
| Negative Edges | 149 | 130 | 242 |
| Average Neighbors | 9.883 | 13.852 | 15.290 |
| Diameter | 24 | 12 | 9 |
| Radius | 12 | 6 | 5 |
| Path Length | 6.474 | 3.652 | 2.887 |
| Clustering Coefficient | 0.478 | 0.481 | 0.481 |
| Density | 0.041 | 0.069 | 0.112 |
| Heterogeneity | 1.064 | 0.926 | 1.021 |
| Centralization | 0.165 | 0.191 | 0.331 |
| Fragmentation | 0.63 | 0.59 | 0.60 |
| Components | 39 | 31 | 33 |

Degree centrality was not significantly different between nearshore and offshore networks (Figure 4A). However, the shelf network had a significantly higher degree centrality than the other two networks. Closeness centrality was significantly different between all three networks (Figure 4B). The offshore network had significantly higher closeness centrality compared to nearshore and shelf. The average shortest path length of nodes was significantly higher in nearshore compared to both shelf and offshore (Figure 4C). Additionally, offshore and shelf had slightly lower fragmentation values (0.60, 0.59) compared to nearshore (0.63) (Table 1).

**Figure 4.**
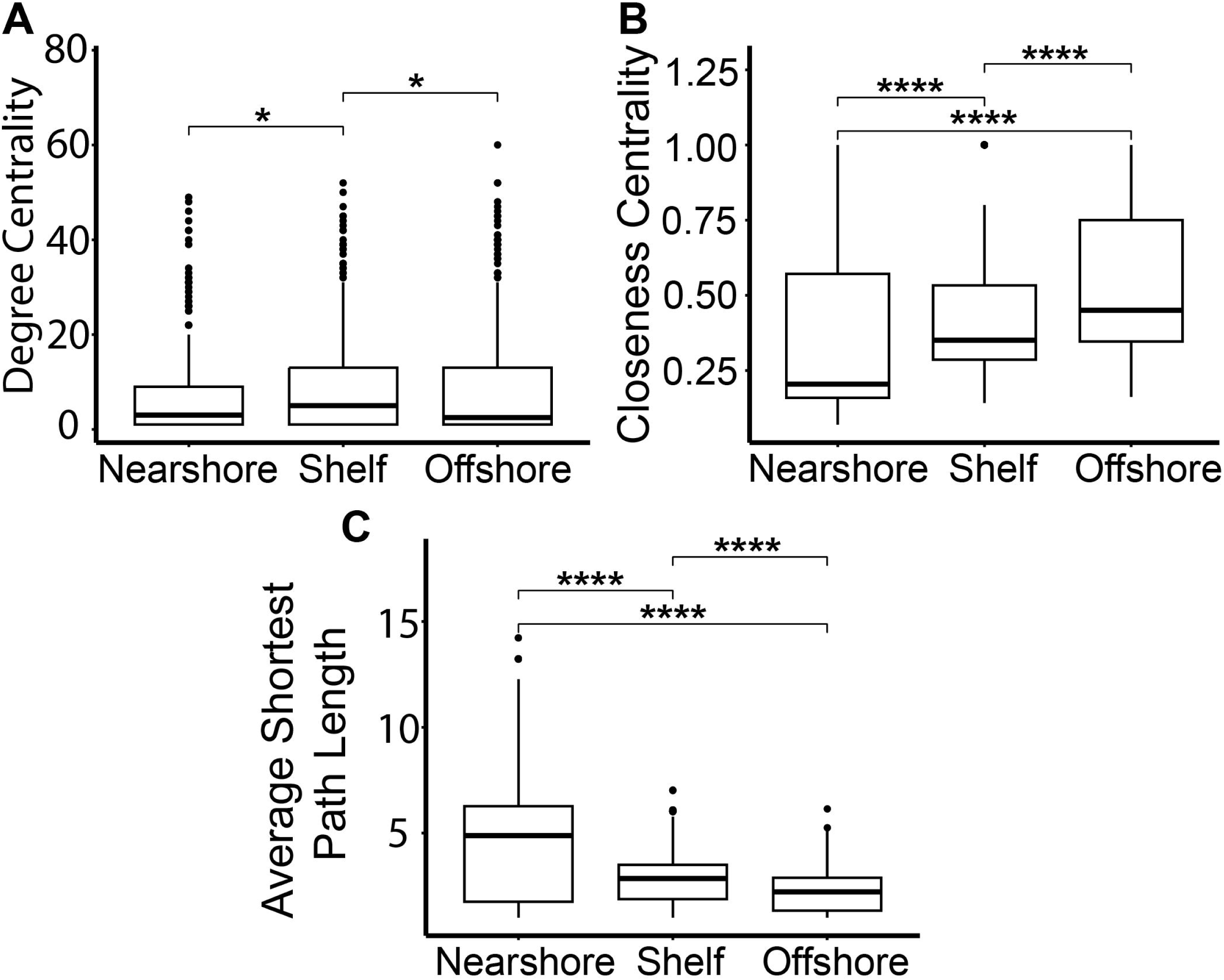
Whole network summary statistics from co-occurrence networks; nearshore, shelf, and offshore. Degree (A), Closeness (B), and Average Shortest Path Length (C) measurements. The number of asterisks represent p-values of Wilcoxon rank sum test (* (*p* ≤ 0.05), ** (*p* ≤ 0.01), *** (*p* ≤ 0.001), **** (*p* ≤ 0.0001)).

### 3.3 Top microbes in network connectivity

Key microbes are those likely to play critical roles within a given network, such that their removal could disrupt the network by severing interactions among microbes. To identify these key microbes within each interaction network, we focused on the top ten microbes with the highest degree centrality, or the most connected nodes in each network (Table 2). These microbes are likely to play critical roles within the networks, such that their removal could disrupt the network by severing the interactions between many microbes. Nearshore and shelf networks were the most similar, having five microbes in common. While nearshore and offshore only had three microbes in common. Only shelf and offshore networks had bacteria in the top ten highest degree centrality. Two of these bacteria were in common, both being within the Proteobacteria phyla including the Gammaproteobacteria SAR86 and the Alphaproteobacteria SAR11 clade IV. Two OTUs were in the top ten in all three networks. Both are within the eukaryote domain: with closest hits to the cryptophyte *P. prolonga* and the haptophyte *Phaeocystis*.

**Table 2.** Potential keystone microbes with the highest degree centrality from each network; Nearshore, Shelf, and Offshore. Closeness centrality is also included.

| Region | Organism | Degree | Closeness |
| --- | --- | --- | --- |
| Nearshore | Eukaryota; *Cryothecomonas aestivalis* | 49 | 0.237 |
| Nearshore | Eukaryota; Dino-Group-I | 48 | 0.237 |
| Nearshore | Eukaryota; MAST-1C | 46 | 0.242 |
| Nearshore | Eukaryota; *Chrysochromulina* sp. | 44 | 0.235 |
| Nearshore | Eukaryota; *Picozoa* sp. | 42 | 0.235 |
| Nearshore | Eukaryota; *Arcocellulus cornucervis* | 42 | 0.234 |
| Nearshore | Eukaryota; *Plagioselmis prolunga* | 42 | 0.234 |
| Nearshore | Eukaryota; Gyrodinium | 40 | 0.239 |
| Nearshore | Eukaryota; Phaeocystis | 39 | 0.232 |
| Nearshore | Eukaryota; *Micromonas commoda* | 34 | 0.237 |
| Shelf | Eukaryota; Dino-Group-I | 52 | 0.433 |
| Shelf | Eukaryota; Phaeocystis | 50 | 0.413 |
| Shelf | Eukaryota; *Plagioselmis prolunga* | 47 | 0.417 |
| Shelf | Bacteria; SAR11 Clade IV | 45 | 0.397 |
| Shelf | Eukaryota; MAST-1C | 45 | 0.406 |
| Shelf | Eukaryota; *Picozoa* sp. | 44 | 0.401 |
| Shelf | Eukaryota; *Bathycoccus prasinos* | 43 | 0.398 |
| Shelf | Bacteria; SAR11 Clade Ia | 42 | 0.402 |
| Shelf | Bacteria; SAR86 | 42 | 0.397 |
| Shelf | Bacteria; NS5 marine group | 40 | 0.392 |
| Offshore | Eukaryota; Gyrodinium | 60 | 0.559 |
| Offshore | Bacteria; AEGEAN-169 | 52 | 0.5 |
| Offshore | Eukaryota; Bacillariophyceae | 52 | 0.515 |
| Offshore | Eukaryota; Phaeocystis | 48 | 0.523 |
| Offshore | Eukaryota; *Plagioselmis prolunga* | 47 | 0.521 |
| Offshore | Bacteria; SAR116 | 46 | 0.491 |
| Offshore | Bacteria; SAR11 Clade IV | 45 | 0.489 |
| Offshore | Eukaryota; Oligotrichida | 45 | 0.498 |
| Offshore | Bacteria; SAR86 | 44 | 0.482 |
| Offshore | Bacteria; SAR86 | 43 | 0.486 |

Microbes with the top ten highest degree centrality in the nearshore network were all eukaryotes (Table 2; Figure 3). Three microbes were only found in the top ten most connected taxa in the nearshore network: the haptophyte *Chrysochromulina* sp., diatom *Arcocellulus cornucervis*, and the green algae *Micromonas commode*. The shelf network, in contrast, had four bacteria in the top ten highest degree centrality OTUs. Two of the four bacteria were Alphaproteobacteria, SAR11 clade IV and SAR11 clade Ia, while the third was a Gammaproteobacteria, SAR86, and the fourth, NS5 marine group differed as a Flavobacteria within the Bacteridota phylum. The offshore network differed the most, having the least common top ten most connected microbes to both nearshore and shelf networks. Bacteria were highest in the offshore network making up half of the ten most connected. All five bacteria were within the Proteobacteria phyla: three Alphaproteobacteria (AEGEAN-169, SAR116, SAR11 clade IV) and two Gammaproteobacteria (SAR86). An unidentified diatom within the Bacillariophyceae class and a ciliate within the Oligotrichida order were also within the top ten most connected microbes in the offshore network.

## 4 Discussion

In the ocean, microbial interactions shape the structure and function of microbial communities, making them critical to marine ecosystem processes (Braga et al., 2016; Arandia-Gorostidi et al., 2022). However, many marine microbes remain uncultivated, and advancing our understanding of their interactions is challenging, particularly because some taxa require co-culturing to survive and grow in laboratory conditions (Lok, 2015; Hofer, 2018; Wang et al., 2021). Network-based analytical approaches such as co-occurrence or co-abundance are a useful alternative, since these can infer potential associations between microbes (Berry and Widder, 2014; Cagua et al., 2019; Yang et al., 2022) and assess ecosystem stability (Widder et al., 2014; Banerjee et al., 2018; Wang et al., 2023; Yang et al., 2024). To uncover potential interactions between microbes in the NGA, we constructed co-occurrence networks of archaea, bacteria, and eukaryotes (Figure 3) and explored network topological features, which allowed us to determine which microbes played an outsized role in the interaction network by being most highly connected (Table 2).

### 4.1 Community composition differed by region

Microbial community composition in both prokaryotes and eukaryotes differed by region in the NGA (Figure 2). Previous research in the tropical Pacific Ocean and Sargasso Sea has observed similar distinct nearshore, shelf, and offshore prokaryotic communities (Wang et al., 2019; Tucker et al., 2021) driven by cross-shelf gradients in salinity and temperature. In the NGA, shifts in prokaryotic and eukaryotic communities were significantly correlated with shifts in salinity and temperature (Figure 2). Prokaryotic communities exhibited interannual variability, consistent with previous observations of community shifts during the 2019 marine heatwave, which favored small celled phytoplankton (Cohen, 2022; Strom, 2023). Other studies in the NGA have reported regional differences in eukaryotic plankton communities, including protists such as dinoflagellates and microzooplankton like ciliates, often linked to cross-shelf salinity gradients and freshwater input from rivers (Coyle and Pinchuk, 2005; Beamer et al., 2016; Strom et al., 2024). We observed a similar trend in our amplicon data, with both prokaryotic and eukaryotic communities clustering by region (Figure 2). Nitrate and phosphate concentrations were highest in offshore and shelf regions and lowest nearshore, indicating stronger macronutrient limitation in surface nearshore waters. This pattern is consistent with prior observations (Strom et al., 2006; Aguilar-Islas et al., 2016; Stabeno et al., 2016; Coyle et al., 2019) and supports the idea that regional niche partitioning of microbial communities is influenced by water mass characteristics.

The ACC has a low-salinity core and transports micronutrients from terrestrial runoff, including inputs from the Copper River plume, to nearshore waters (Ortega et al., 2025). Although rich in micronutrients like iron and copper, the ACC is often limited by macronutrients such as nitrate during summer (Strom et al., 2006; Aguilar-Islas et al., 2016; Coyle et al., 2019). Past studies have observed chain-forming diatoms dominating these nearshore waters, particularly during summer stratification (Strom et al., 2006). Our amplicon data supported this pattern, with high relative read abundance of Bacillariophyceae diatoms, including *Pseudo-nitzschia* spp. and *Arcocellulus* spp., in nearshore samples (Figure S1E). *Pseudo-nitzschia* is a chain-forming pennate diatom frequently associated with elevated nutrient concentrations and high productivity (Zhang et al., 2021; Moreno et al., 2022), traits characteristic of the ACC (Strom et al., 2006; Stabeno et al., 2016). Some *Pseudo-nitzschia* species are also known to produce domoic acid, a neurotoxin that causes amnesic shellfish poisoning; however, surveys in the NGA to date have not detected domoic acid levels exceeding regulatory limits (Alaska Harmful Algal Bloom Network, 2023). In contrast, offshore waters showed high relative abundance of *Actinocyclus*, a large centric diatom. While often associated with nutrient-rich coastal regions, *Actinocyclus* can also appear in offshore waters when mesoscale eddies or cross-shelf exchange events introduce nutrient pulses to otherwise iron-limited surface waters (Stabeno et al., 2004; Crawford et al., 2007; Ladd et al., 2007). This regional distinction in diatom assemblages likely reflects niche partitioning shaped by nutrient availability along cross-shelf gradients and water masses.

### 4.2 Co-occurrence network suggests the importance of kleptoplasty and parasitoids in NGA

To examine microbial interactions shared across environmental gradients, we constructed a merged network of significant correlations and nodes found in all three regions. Co-occurrence networks infer potential interactions based on patterns of co-presence or mutual exclusion, but these patterns can result from various ecological processes, including direct interactions, shared environmental preferences, or indirect associations (Fuhrman, 2009; Weiss et al., 2016). Positive correlations may indicate mutualism, metabolic exchange, predation, or antagonistic exclusion (Hibbing et al., 2010; Ghoul and Mitri, 2016). However, correlation alone does not imply causation, and method limitations such as the lack of temporal resolution and potential indirect effects must be considered when interpreting these networks (Needham et al., 2013; Kodera et al., 2022). Despite these constraints, co-occurrence networks are useful hypothesis-generating tools that can identify recurring ecological patterns and guide future experimental investigations.

The merged network revealed significant correlations and nodes shared across all three regions. Although there were significant differences between nearshore and offshore networks and different potential keystone microbes, some significant correlations were found in all three networks (Figure 5). The cryptophyte *P. prolonga* was a potential keystone microbe in all three networks (Figure 3; Table 2). *P. prolonga* is a common marine cryptophyte that is an important plastid donor in kleptoplastic interactions with ciliates and dinoflagellates (Altenburger et al., 2020; Cruz and Cartaxana, 2022). In our networks, *P. prolonga* was consistently associated with ciliates and dinoflagellates, including a grouping found in all three regions involving the ciliate order Oligotrichida and the dinoflagellate *Gyrodinium* (Figures 5 and 7). Kleptopasty allows the host to obtain plastids from ingested photosynthetic prey (Tsuchiya et al., 2020; Cruz and Cartaxana, 2022). Marine oligotrich ciliates have been found to retain prey plastids which remain functional for days to weeks giving them the temporary ability to perform photosynthesis in addition to phagotrophy, making them non-constitutive mixotrophs (Stoecker et al., 1988; Jonsson, 1989; McManus et al., 2012). Ciliates within Oligotrichida have been identified in high latitude environments with the ability to perform mixotrophy (Stoecker and Lavrentyev, 2018; O’Hara, 2023; Strom et al., 2024), suggesting this strategy may be especially advantageous in cold, seasonally variable systems like the NGA. Mixotrophy enhances survival under low-prey or low-light conditions and supports vertical and horizontal transfer of organic matter (Stoecker et al., 2017). It can also lengthen food chains and improve trophic transfer efficiency by bridging microbial and metazoan consumers (Stoecker et al., 2017). Recent work highlights the ecological significance of mixotrophs in the NGA, where environmental gradients in light and nutrients may select for flexible nutritional modes (Strom et al., 2024). The consistent correlations between *P. prolonga* and Oligotrichida nodes suggest kleptoplasty is a prevalent nutritional strategy in the NGA, contributing to primary production and enhancing trophic connectivity in this high-latitude ecosystem.

**Figure 5.**
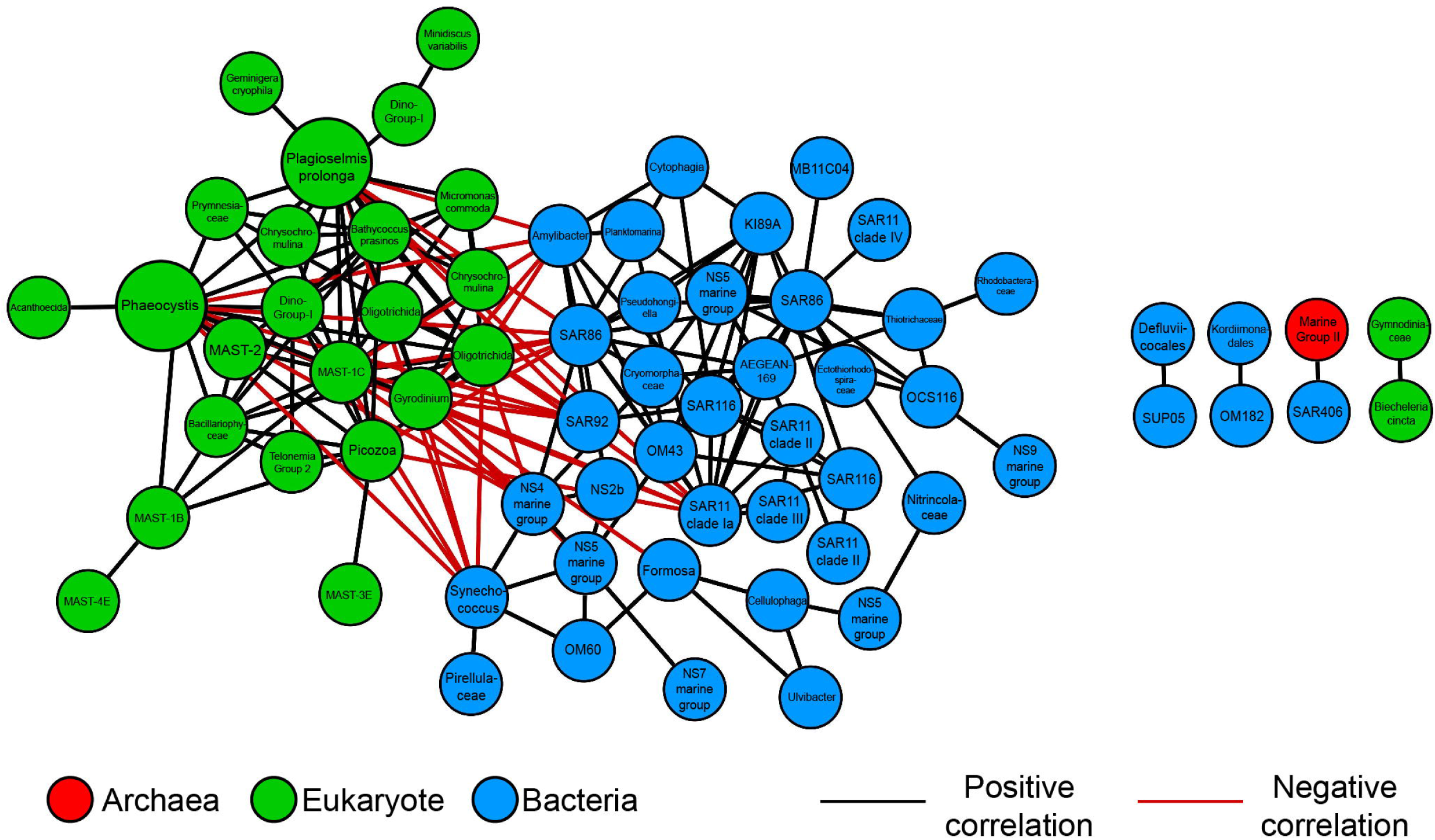
Merged network of significant correlations and nodes found in all three networks (Nearshore, Shelf, and Offshore). Black edges represent positive significant correlations, while red represents negative correlations. Nodes are color-coded by taxonomy: Archaea (red), Bacteria (blue), and Eukaryota (green). Larger nodes are potential keystone microbes from global networks.

Within the nearshore network microbes associated with parasitism and predation were more abundant and connected (Figure 3). This includes potential keystone microbes and predatory BALOs (Figure 6; Table 2). BALOs are parasitoid bacteria that feed upon and reproduce within gram-negative bacteria using bdelloplasts until the prey cell lyses releasing new BALO cells (Sockett, 2009; Negus et al., 2017; Laloux, 2019). BALOs have two life phases a motile attack phase where they can swim toward prey and a proliferative phase after attachment to prey (Sockett, 2009; Laloux, 2019). Subnetworks showed BALOs (including OM27) significantly correlated with flavobacteria such as NS7 marine group, Cryomorphaceae, and others in all three regions (Figure 6). These flavobacteria are often particle-associated marine microbes (Mitulla et al., 2016; Milici et al., 2017). Since BALOs contain two different life phases, they may move toward particles where there is a higher abundance of potential prey (Lambert et al., 2006, 2011; Iida et al., 2009). Flow cytometry data showed higher heterotrophic bacteria and *Synechococcus* nearshore cell counts compared to offshore (Figure S2). Prey density has been linked to predator density as well as predator-prey interactions (Holling, 1959; Dick et al., 2014; Cuthbert et al., 2021). A minimum prey density is therefore required to support a predator population (Holling, 1959; Dick et al., 2014; Cuthbert et al., 2021). Higher cell counts could increase cell contact rates and therefore give parasitic and predatory microbes higher success in the nearshore network (Hu et al., 2013).

**Figure 6.**
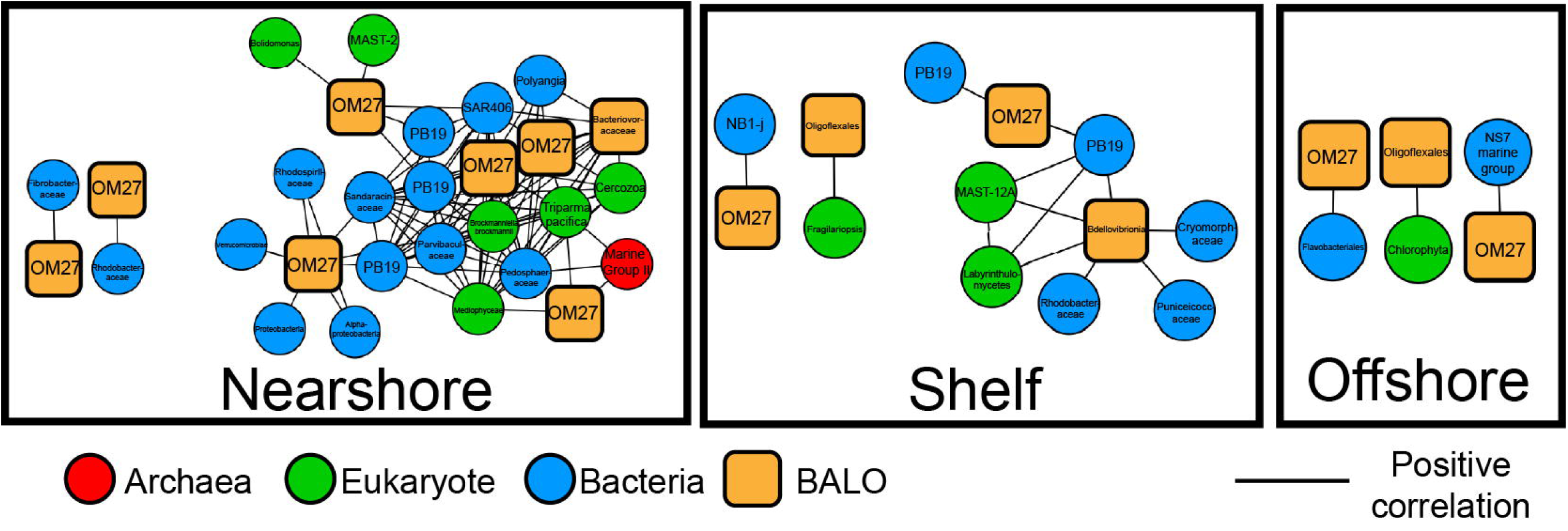
Subnetwork of predatory Bdellovibrio and like organisms (BALOs) (orange squares) and significantly correlated nodes, including potential bacterial prey (blue circles) and associations with eukaryotes (green circles).

**Figure 7.**
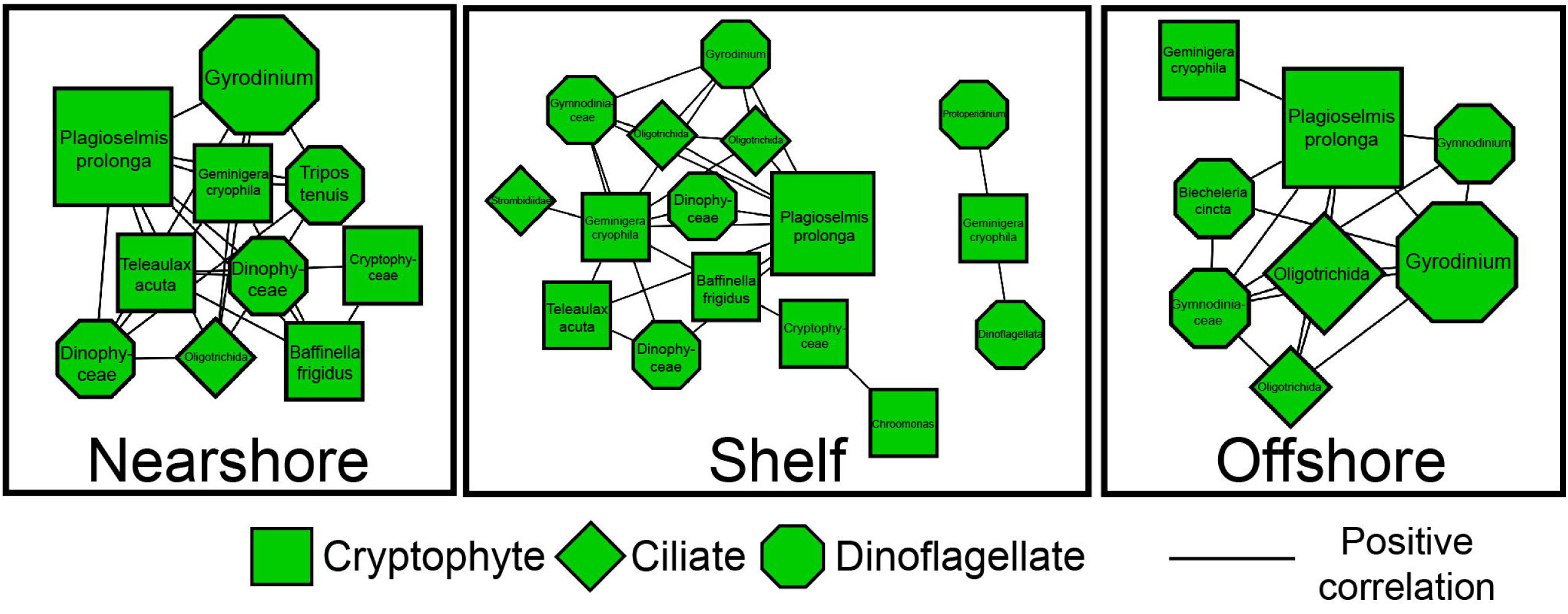
Subnetworks of cryptophyte species (squares) and significantly correlated ciliate (diamond) and dinoflagellate (octagon) nodes. Larger nodes are potential keystone microbes from global networks.

Interestingly, eukaryotic parasitoids, such as *Cryothecomonas aesticalis*, a nanoflagellate parasitizing diatoms (Drebes et al., 1996; Catlett et al., 2023), were the most connected microbes in the nearshore network (Table 2), highlighting how complex microbial interactions like parasitism can dominate under dynamic, high-biomass conditions even when surface nutrients are seasonally depleted (Worden et al., 2015). Other highly connected microbes in the nearshore, such as Syndiniales Dino-Group I and MAST-1C, are also known or suspected parasites (Guillou et al., 2008; Massana et al., 2009; Lin et al., 2012; Cleary and Durbin, 2016; Clarke et al., 2019). These eukaryotic parasites were notably absent as the most connected microbes in the offshore network, where microbial parasites were not considered keystone species. The nearshore environment typically has higher micronutrient availability; however, in summer, surface macronutrient depletion may limit primary production, allowing excess micronutrients to persist. These conditions could still support abundant microbial communities particularly in subsurface layers or particle-rich zones that serve as hosts for parasites (Mitulla et al., 2016; Milici et al., 2017). Additionally, freshwater inputs nearshore often create a shallow pycnocline, which may concentrate potential hosts in the surface layer and enhance contact rates with parasites (Lucas et al., 1999). Greater environmental variability in salinity and temperature nearshore may also increase host susceptibility to infection, as physiological stress can compromise host defenses (Harvell et al., 2002). In addition, in macronutrient-depleted surface waters, some microbes may occupy deeper layers to access nutrients, where reduced light imposes further metabolic stress that could increase vulnerability to parasitism (Strom et al., 2006; Worden et al., 2015; Stoecker et al., 2017). In contrast, offshore waters are typically colder and iron-limited, with high nitrate but low chlorophyll concentrations characteristic of high-nutrient, low-chlorophyll (HNLC) systems (Aguilar-Islas et al., 2016; Coyle et al., 2019). Host density in these waters may be limited by iron availability, reducing opportunities for parasitic lifestyles (Boyd, 2007; Strom et al., 2024). However, episodic mixing events such as eddies or intrusions from the Alaska Current and Alaskan Stream may intermittently deliver iron to surface waters, potentially creating localized conditions favorable for parasitism (Ladd et al., 2007; Coyle et al., 2019). These ecological factors may help explain the stronger prominence of parasitic eukaryotes in the nearshore compared to offshore ecosystems.

### 4.3 What do network characteristics suggest about resilience and stability of the NGA?

Network analysis of microbial co-occurrences provides valuable insights into community structure, resilience, and stability by identifying keystone taxa, those microbes most crucial to maintaining ecological functions. Using measures like degree centrality, which reflects a taxon’s connectivity within the network, we can determine which species, if removed, would be most likely to disrupt community dynamics (Berry and Widder, 2014; Banerjee et al., 2018). The loss of these keystone microbes can lead to significant shifts in microbial community structure and function, as their absence can disproportionately affect ecological processes (Berry and Widder, 2014; Cagua et al., 2019). Network fragmentation, which occurs when key correlations between taxa are lost, further exacerbates this disruption, leading to communities that are less resilient to disturbances (Widder et al., 2014; Wang et al., 2023; Yang et al., 2024). Communities exhibiting higher functional redundancy and lower fragmentation are more likely to maintain stability and recover from perturbations, as they have a greater capacity to compensate for the loss of keystone taxa (Wang et al., 2023). Understanding these key interactions and network properties is essential when evaluating the stability and adaptability of microbial communities, especially in varying environmental conditions.

Potential keystone microbes were identified as the top ten taxa in each network with the highest degree centrality values (Table 2; Figure 3). Our results suggest that potential keystone microbes shifted from predominantly eukaryotic taxa nearshore to an equal mix of bacteria and eukaryotes offshore (Table 2; Figure 3). Offshore keystone microbes included many oligotrophic bacteria such as SAR11, SAR86, and SAR116 (Brown et al., 2012; Giovannoni, 2017; Hoarfrost et al., 2020; Roda-Garcia et al., 2021). While some of these taxa are also highly abundant, which may contribute to their high centrality, their consistent appearance as central nodes across samples suggests they may also play important ecological roles. Despite being abundant and widespread, many bacteria with oligotrophic lifestyles are adapted to low-nutrient conditions and remain poorly understood due to their uncultivated status (Faust and Raes, 2012; Zelezniak et al., 2015; Yan et al., 2023). These traits may confer an advantage in the iron-limited HNLC offshore waters of the NGA. Studies have shown that interactions among such streamlined taxa, including cross-feeding and metabolite exchange, can structure microbial communities and enhance ecosystem function, highlighting the importance of further investigating their ecological roles (Braakman et al., 2017).

The negative correlations between bacteria and eukaryotic keystone microbes found in our datasets (Figure 3) are possibly driven by bacterivory, with eukaryotic microbes phagocytizing bacterial prey (Sakka et al., 2000; Christaki et al., 2002; Strom, 2002; Sarmento et al., 2013; Arandia-Gorostidi et al., 2022; Follett et al., 2022). For example, the cryptophyte *Plagioselmis prolonga*, a keystone microbe in the nearshore network, had exclusively negative correlations with bacteria (Figure 5). Many cryptophytes are mixotrophs, supplementing autotrophy with phagotrophy of bacteria (O’Hara, 2023; Šimek et al., 2023), suggesting that *P. prolonga’s* role as a keystone microbe may be linked to its dual trophic strategy. Similarly, *Phaeocystis*, another keystone microbe present across all regions, had a high number of negative correlations with bacteria, possibly due to bacterial degradation of bloom remains, such as carbon released from cell lysis (Brussaard et al., 1995, 1996) or due to the ability of *Phaeocystis* to phagocytize bacteria (Koppelle et al., 2022). By linking microbial primary production and heterotrophic consumption, mixotrophy may enhance microbial connectivity and ecosystem function, reinforcing the keystone roles of *P. prolonga* and *Phaeocystis* (Li et al., 2024).

Network-wide stability metrics indicated that the offshore network exhibited nodes with significantly higher closeness centrality (Figure 4B) compared to the networks in the other two regions. Higher closeness centrality suggests that microbes in the offshore network are more interconnected, meaning disturbances may propagate more quickly, but the overall network is more resistant to fragmentation (Assenov et al., 2008; Berry and Widder, 2014; Cagua et al., 2019; Matchado et al., 2021). While we cannot directly infer specific interaction types from correlation-based networks, higher connectivity in other ecological systems has been linked to community stability and robustness (Jeong et al., 2000; Poisot et al., 2011; Hannigan et al., 2018; Qian and Akçay, 2020). Some studies have also found that highly connected networks are more common in systems where cooperative or mutually beneficial interactions prevail (Thébault and Fontaine, 2010; Montesinos-Navarro et al., 2017; Costas-Selas et al., 2024). Further investigation is needed to confirm whether similar dynamics are occurring in the NGA. In contrast, lower connectivity in the nearshore and shelf networks could indicate reduced stability, potentially driven by a greater prevalence of antagonistic interactions such as parasitism or competition. Alternatively, it may reflect greater environmental variability in these regions than offshore, which could disrupt consistent associations among microbes and lead to higher fragmentation and susceptibility to disturbance. Taken together, these patterns suggest that while the offshore network may be more resistant to perturbation due to its higher connectivity, the lower connectivity observed in nearshore and shelf communities could either make them more vulnerable to environmental change or allow for greater adaptability in response to the higher variability they experience.

Defining a stable ecosystem, especially a marine one, is difficult since the system is inherently dynamic. Researchers have used fragmentation measurements of microbial co-occurrence networks as an indicator of ecological network sensitivity, with higher fragmentation suggesting greater instability (Widder et al., 2014; Banerjee et al., 2018; Yang et al., 2024). All three co-occurrence networks had fragmentation values greater than 0.5, suggesting microbial communities in all three regions of the NGA are more fragmented and potentially less stable than those observed in other marine ecosystems with lower fragmentation, such as freshwater lates, the East China Sea, and seagrass meadows (Widder et al., 2014; Wang et al., 2023; Yang et al., 2024). Among the three regions, the nearshore network had the highest fragmentation value and the lowest connectivity (Table 1), which could indicate greater vulnerability to disturbance. This region also experiences the highest environmental variability, such fragmentation may also reflect greater flexibility in microbial associations rather than instability per se. This region also contained a greater proportion of parasitic and predatory taxa, such as parasitic eukaryotes and BALOs, which have been shown to influence microbial community structure through top-down interactions and host-parasite dynamics (Thingstad and Lignell, 1997; Chow et al., 2013; Lima-Mendez et al., 2015; Worden et al., 2015). The presence of these taxa suggests strong top-down control, which may contribute to increased fragmentation. According to the kill-the-winner hypothesis, strain-specific predators like viruses can promote microbial diversity by targeting the most competitive taxa, preventing dominance and enabling coexistence (Thingstad, 2000; Winter et al., 2010; Thingstad et al., 2014). If parasitism disproportionately affects highly connected or central taxa, it may destabilize the network by weakening key interactions (Weitz and Dushoff, 2008; Chow et al., 2013; Lima-Mendez et al., 2015). Alternatively, top down pressure could allow less dominant taxa to emerge as new keystone species, potentially reshaping the structure and function of the microbial community. The high fragmentation in the nearshore network could reflect this where top-down control both supports diversity and increases the risk of instability.

Differences in network characteristics between nearshore and offshore regions may be driven by two key factors. First, microbial community composition and keystone taxa vary significantly between regions (Figure 2; Table 2), influencing interaction types and network structure. Antagonistic interactions, such as parasitism or predation, may be more prevalent in the nearshore, where exploitative relations could dominate, whereas offshore microbial communities may experience a more complex mix of interactions. These interactions might not necessarily be mutualistic but could involve processes like micronutrient sharing or cross-feeding that may foster greater network cohesion (Amin et al., 2012; Bertrand et al., 2015; Cooper and Smith, 2015; Arandia-Gorostidi et al., 2022). Second, geochemical differences between regions, driven by the mixing of water masses in the Northern Gulf of Alaska, impact nutrient availability and primary productivity (Strom et al., 2010; Aguilar-Islas et al., 2016; Coyle et al., 2019). Offshore microbial communities may be more stable, potentially due to selective pressures favoring micronutrient exchange processes such as cross-feeding or siderophore exchange, which could reinforce network stability (Amin et al., 2012; Morris et al., 2012; Zelezniak et al., 2015; D’Souza et al., 2018). In contrast, the nearshore environment, characterized by more variable micronutrient and freshwater inputs, may foster a greater prevalence of exploitative interactions. The differences suggest that moderate variability in nutrient availability and a mosaic environmental structure could promote microbial diversity and stability, but this remains speculative and contrasts with the observed fragmentation in the nearshore network. Ultimately, these findings raise questions about the potential for ecosystem transitions in the NGA. In 2019, a shift in microbial community composition was observed, including a notable increase in chlorophytes and oligotrophic bacteria (Cohen, 2022). The challenge moving forward is to identify early warning signs that might precede tipping points in ecosystem functioning. Identifying these signs could help predict future transitions and offer valuable insights for understanding the resilience of the system (Chisholm and Filotas, 2009; Scheffer et al., 2012).

### 4.4 Conclusions

This study found microbial communities, potential keystone microbes, and their correlations differ across the shelf/slope of the NGA. Potential keystone microbes that were the most connected in the micronutrient-rich nearshore network were all eukaryotes. However, as distance increased to offshore waters, there was a shift in keystone microbes from eukaryotes to smaller bacteria often found in oligotrophic waters. The offshore network exhibited higher connectivity and less fragmentation compared to the nearshore, suggesting it may be more stable either due to greater microbial interconnectivity or as a result of the less variable environmental conditions offshore. This stability may be further promoted by more cross-domain interactions and a more balanced representation between bacteria and eukaryotes as keystone microbes in the offshore network. While the nearshore network seemed to have a greater prevalence of possible antagonistic interactions, the offshore network likely involves a more complex mix of interactions, with processes like micronutrient sharing or cross-feeding potentially playing a role in fostering network cohesion. The difference in potential interactions suggests a shift from more exploitative relationships nearshore to a more cooperative system offshore where low micronutrient concentrations can be an environmental stressor. Additionally, this study identified two potential key microbial players found in all regions, the mixotrophic nanoflagellates *Plagioselmis prolonga* and *Phaeocystis* sp., suggesting these microbes strategies are well suited to environmental gradients across the shelf/slope domains, and may contribute disproportionately to microbial interaction network stability. Overall, different network characteristics and potential interactions were apparent by region, but further research is needed to determine how microbial community composition and environmental variability influence ecosystem stability. These findings contribute to the understanding of microbial ecology during summer conditions in the NGA, and could have important implications for understanding the resilience or vulnerability of the NGA in the face of climate change.

## Acknowledgements

We thank the captains and crew of the *R/V Sikuliaq* and *R/V Woldstad*, Dr. Suzanne Strom and the members of the Strom lab, Dr. Ana Aguilar-Islas, and the UAF Water Nutrient Analytical Facility. This work was supported by funding from NSF Biological Oceanography (OCE 1851101), the Northern Gulf of Alaska Long Term Ecological Research site (OCE 1656070), and NSF Office of Polar Programs (OPP 1937595).

## Author Contributions

MB led the project conceptualization, performed all data curation, conducted bioinformatic and statistical analyses, and wrote the original draft. JC collected and processed field and molecular samples. BRB contributed to project conceptualization, provided guidance on the bioinformatic workflow, assisted with manuscript review and editing, and contributed to software development. GMMH contributed to project conceptualization, supervised the research, participated in sample collection, coordinated sample processing logistics, provided critical input as a lead investigator in the NGA LTER program, and contributed to funding acquisition. Both BRB and GMMH were involved in acquiring financial support for the project. All authors contributed to the review and editing of the final manuscript and approved the submitted version.

## Data Availability Statement

Demultiplexed sequencing data for this study have been deposited in the NCBI Sequence Read Archive under BioProject accession number PRJNA887083. Individual samples are available under BioSample accession numbers SAMN31684678–SAMN31685019.

## References

Abirami, B., Radhakrishnan, M., Kumaran, S., and Wilson, A. (2021). Impacts of global warming on marine microbial communities. Sci Total Environ 791, 147905. doi: 10.1016/j.scitotenv.2021.147905

Aguilar-Islas, A. M., Séguret, M. J. M., Rember, R., Buck, K. N., Proctor, P., Mordy, C. W., et al. (2016). Temporal variability of reactive iron over the Gulf of Alaska shelf. Deep Sea Research Part II: Topical Studies in Oceanography 132, 90–106. doi: 10.1016/j.dsr2.2015.05.004

Alaska Harmful Algal Bloom Network (2023). 2022 Alaska Summary Report of Regional Harmful Algal Bloom Activities. Alaska Harmful Algal Bloom Network. Available at: https://ahab.aoos.org/wp-content/uploads/2023/10/AHAB-2022-Summary-Report-final.pdf

Altenburger, A., Blossom, H. E., Garcia-Cuetos, L., Jakobsen, H. H., Carstensen, J., Lundholm, N., et al. (2020). Dimorphism in cryptophytes—The case of Teleaulax amphioxeia/Plagioselmis prolonga and its ecological implications. Science Advances 6, eabb1611. doi: 10.1126/sciadv.abb1611

Amaral-Zettler, L. A., McCliment, E. A., Ducklow, H. W., and Huse, S. M. (2009). A Method for Studying Protistan Diversity Using Massively Parallel Sequencing of V9 Hypervariable Regions of Small-Subunit Ribosomal RNA Genes. PLOS ONE 4, e6372. doi: 10.1371/journal.pone.0006372

Amin, S. A., Parker, M. S., and Armbrust, E. V. (2012). Interactions between Diatoms and Bacteria. Microbiol Mol Biol Rev 76, 667–684. doi: 10.1128/MMBR.00007-12

Arandia-Gorostidi, N., Krabberød, A. K., Logares, R., Deutschmann, I. M., Scharek, R., Morán, X. A. G., et al. (2022). Novel Interactions Between Phytoplankton and Bacteria Shape Microbial Seasonal Dynamics in Coastal Ocean Waters. Front. Mar. Sci. 9, 901201. doi: 10.3389/fmars.2022.901201

Assenov, Y., Ramírez, F., Schelhorn, S.-E., Lengauer, T., and Albrecht, M. (2008). Computing topological parameters of biological networks. Bioinformatics 24, 282–284. doi: 10.1093/bioinformatics/btm554

Banerjee, S., Schlaeppi, K., and van der Heijden, M. G. A. (2018). Keystone taxa as drivers of microbiome structure and functioning. Nature Reviews Microbiology 16, 567–576. doi: 10.1038/s41579-018-0024-1

Basu, S., Duren, W., Evans, C. R., Burant, C. F., Michailidis, G., and Karnovsky, A. (2017). Sparse network modeling and metscape-based visualization methods for the analysis of large-scale metabolomics data. Bioinformatics 33, 1545–1553. doi: 10.1093/bioinformatics/btx012

Beamer, J. P., Hill, D. F., Arendt, A., and Liston, G. E. (2016). High-resolution modeling of coastal freshwater discharge and glacier mass balance in the Gulf of Alaska watershed. Water Resources Research 52, 3888–3909. doi: 10.1002/2015WR018457

Behrenfeld, M. J., O’Malley, R. T., Siegel, D. A., McClain, C. R., Sarmiento, J. L., Feldman, G. C., et al. (2006). Climate-driven trends in contemporary ocean productivity. Nature 444, 752–755. doi: 10.1038/nature05317

Berry, D., and Widder, S. (2014). Deciphering microbial interactions and detecting keystone species with co-occurrence networks. Frontiers in Microbiology 5. doi: 10.3389/fmicb.2014.00219

Bertrand, E. M., McCrow, J. P., Moustafa, A., Zheng, H., McQuaid, J. B., Delmont, T. O., et al. (2015). Phytoplankton–bacterial interactions mediate micronutrient colimitation at the coastal Antarctic sea ice edge. PNAS 112, 6.

Bodor, A., Bounedjoum, N., Vincze, G. E., Erdeiné Kis, Á., Laczi, K., Bende, G., et al. (2020). Challenges of unculturable bacteria: environmental perspectives. Rev Environ Sci Biotechnol 19, 1–22. doi: 10.1007/s11157-020-09522-4

Bokulich, N. A. (2018). Optimizing taxonomic classification of marker-gene amplicon sequences with QIIME 2’s q2-feature-classifier plugin. 17.

Bolyen, E., Rideout, J. R., Dillon, M. R., Bokulich, N. A., Abnet, C. C., Al-Ghalith, G. A., et al. (2019). Reproducible, interactive, scalable and extensible microbiome data science using QIIME 2. Nat Biotechnol 37, 852–857. doi: 10.1038/s41587-019-0209-9

Boyd, P. W. (2007). Biogeochemistry: iron findings. Nature 446, 989–91. doi: 10.1038/446989a

Boyd, P. W., Doney, S. C., Strzepek, R., Dusenberry, J., Lindsay, K., and Fung, I. (2008). Climate-mediated changes to mixed-layer properties in the Southern Ocean: assessing the phytoplankton response. Biogeosciences 5, 847–864. doi: 10.5194/bg-5-847-2008

Braakman, R., Follows, M. J., and Chisholm, S. W. (2017). Metabolic evolution and the self-organization of ecosystems. Proceedings of the National Academy of Sciences 114, E3091–E3100. doi: 10.1073/pnas.1619573114

Braga, R. M., Dourado, M. N., and Araújo, W. L. (2016). Microbial interactions: ecology in a molecular perspective. Brazilian Journal of Microbiology 47, 86–98. doi: 10.1016/j.bjm.2016.10.005

Brown, M. V., Lauro, F. M., DeMaere, M. Z., Muir, L., Wilkins, D., Thomas, T., et al. (2012). Global biogeography of SAR11 marine bacteria. Mol Syst Biol 8, 595. doi: 10.1038/msb.2012.28

Brussaard, C., Gast, G. J., Duyl, F., and Riegman, R. (1996). Impact of phytoplankton bloom magnitude on a pelagic microbial food web. Marine Ecology Progress Series 144, 211–221. doi: 10.3354/meps144211

Brussaard, C., Riegman, R., Noordeloos, A., Cadee, G., Witte, H., Kop, A. J., et al. (1995). Effects of grazing, sedimentation and phytoplankton cell lysis on the structure of a coastal pelagic food web. Marine Ecology Progress Series 123, 259–271. doi: 10.3354/meps123259

Cagua, E. F., Wootton, K. L., and Stouffer, D. B. (2019). Keystoneness, centrality, and the structural controllability of ecological networks. Journal of Ecology 107, 1779–1790. doi: 10.1111/1365-2745.13147

Callahan, B. J., McMurdie, P. J., Rosen, M. J., Han, A. W., Johnson, A. J. A., and Holmes, S. P. (2016). DADA2: High-resolution sample inference from Illumina amplicon data. Nat Methods 13, 581–583. doi: 10.1038/nmeth.3869

Catlett, D., Peacock, E., Crockford, E., Futrelle, J., Batchelder, S., Stevens, B., et al. (2023). Temperature dependence of parasitoid infection and abundance of a diatom revealed by automated imaging and classification. Proceedings of the National Academy of Sciences of the United States of America 120, e2303356120. doi: 10.1073/pnas.2303356120

Chisholm, R. A., and Filotas, E. (2009). Critical slowing down as an indicator of transitions in two-species models. Journal of Theoretical Biology 257, 142–149. doi: 10.1016/j.jtbi.2008.11.008

Chow, C.-E. T., Sachdeva, R., Cram, J. A., Steele, J. A., Needham, D. M., Patel, A., et al. (2013). Temporal variability and coherence of euphotic zone bacterial communities over a decade in the Southern California Bight. ISME J 7, 2259–2273. doi: 10.1038/ismej.2013.122

Christaki, U., Courties, C., Karayanni, H., Giannakourou, A., Maravelias, C., Kormas, K. Ar., et al. (2002). Dynamic Characteristics of Prochlorococcus and Synechococcus Consumption by Bacterivorous Nanoflagellates. Microbial Ecology 43, 341–352.

Clarke, L. J., Bestley, S., Bissett, A., and Deagle, B. E. (2019). A globally distributed Syndiniales parasite dominates the Southern Ocean micro-eukaryote community near the sea-ice edge. ISME J 13, 734–737. doi: 10.1038/s41396-018-0306-7

Cleary, A. C., and Durbin, E. G. (2016). Unexpected prevalence of parasite 18S rDNA sequences in winter among Antarctic marine protists. Journal of Plankton Research 38, 401–417. doi: 10.1093/plankt/fbw005

Cock, P. J. A., Antao, T., Chang, J. T., Chapman, B. A., Cox, C. J., Dalke, A., et al. (2009). Biopython: freely available Python tools for computational molecular biology and bioinformatics. Bioinformatics 25, 1422–1423. doi: 10.1093/bioinformatics/btp163

Cohen, J. (2022). Shifts in microbial community composition during the 2019 Pacific marine heatwave in the northern Gulf of Alaska. Available at: http://hdl.handle.net/11122/13115

Cohen, J., Screen, J. A., Furtado, J. C., Barlow, M., Whittleston, D., Coumou, D., et al. (2014). Recent Arctic amplification and extreme mid-latitude weather. Nature Geoscience 7, 627–637. doi: 10.1038/ngeo2234

Cohen, Y., Pasternak, Z., Müller, S., Hübschmann, T., Schattenberg, F., Sivakala, K. K., et al. (2021). Community and single cell analyses reveal complex predatory interactions between bacteria in high diversity systems. Nature Communications 12, 5481. doi: 10.1038/s41467-021-25824-9

Cooper, M. B., and Smith, A. G. (2015). Exploring mutualistic interactions between microalgae and bacteria in the omics age. Current Opinion in Plant Biology 26, 147–153. doi: 10.1016/j.pbi.2015.07.003

Costas-Selas, C., Martínez-García, S., Delgadillo-Nuño, E., Justel-Díez, M., Fuentes-Lema, A., Fernández, E., et al. (2024). Linking the impact of bacteria on phytoplankton growth with microbial community composition and co-occurrence patterns. Marine Environmental Research 193, 106262. doi: 10.1016/j.marenvres.2023.106262

Coyle, K. O., Hermann, A. J., and Hopcroft, R. R. (2019). Modeled spatial-temporal distribution of productivity, chlorophyll, iron and nitrate on the northern Gulf of Alaska shelf relative to field observations. Deep Sea Research Part II: Topical Studies in Oceanography 165, 163–191. doi: 10.1016/j.dsr2.2019.05.006

Coyle, K. O., and Pinchuk, A. I. (2005). Seasonal cross-shelf distribution of major zooplankton taxa on the northern Gulf of Alaska shelf relative to water mass properties, species depth preferences and vertical migration behavior. Deep Sea Research Part II: Topical Studies in Oceanography 52, 217–245. doi: 10.1016/j.dsr2.2004.09.025

Cram, J. A., Chow, C.-E. T., Sachdeva, R., Needham, D. M., Parada, A. E., Steele, J. A., et al. (2015). Seasonal and interannual variability of the marine bacterioplankton community throughout the water column over ten years. The ISME Journal 9, 563–580. doi: 10.1038/ismej.2014.153

Crawford, W., Brickley, P., and Thomas, A. (2007). Mesoscale eddies dominate surface phytoplankton in northern Gulf of Alaska. Progress in Oceanography - PROG OCEANOGR 75. doi: 10.1016/j.pocean.2007.08.016

Cruz, S., and Cartaxana, P. (2022). Kleptoplasty: Getting away with stolen chloroplasts. PLoS Biol 20, e3001857. doi: 10.1371/journal.pbio.3001857

Cui, Y., Chun, S.-J., Baek, S. H., Lee, M., Kim, Y., Lee, H.-G., et al. (2019). The water depth-dependent co-occurrence patterns of marine bacteria in shallow and dynamic Southern Coast, Korea. Sci Rep 9, 9176. doi: 10.1038/s41598-019-45512-5

Cuthbert, R. N., Dalu, T., Wasserman, R. J., Sentis, A., Weyl, O. L. F., Froneman, P. W., et al. (2021). Prey and predator density-dependent interactions under different water volumes. Ecol Evol 11, 6504–6512. doi: 10.1002/ece3.7503

Dick, J. T. A., Alexander, M. E., Jeschke, J. M., Ricciardi, A., MacIsaac, H. J., Robinson, T. B., et al. (2014). Advancing impact prediction and hypothesis testing in invasion ecology using a comparative functional response approach. Biological Invasions 16, 735–753. doi: 10.1007/s10530-013-0550-8

Diestel, R. (2005). Graph Theory. Springer New York.

Dixon, P. (2003). VEGAN, a package of R functions for community ecology. Journal of Vegetation Science 14, 927–930. doi: 10.1111/j.1654-1103.2003.tb02228.x

Drebes, G., Kühn, S. F., Gmelch, A., and Schnepf, E. (1996). Cryothecomonas aestivalis sp. nov., a colourless nanoflagellate feeding on the marine centric diatomGuinardia delicatula (Cleve) Hasle. Helgoländer Meeresuntersuchungen 50, 497–515.

D’Souza, G., Shitut, S., Preussger, D., Yousif, G., Waschina, S., and Kost, C. (2018). Ecology and evolution of metabolic cross-feeding interactions in bacteria. Nat. Prod. Rep. 35, 455–488. doi: 10.1039/C8NP00009C

Faust, K., and Raes, J. (2012). Microbial interactions: from networks to models. Nature Reviews Microbiology 10, 538–550. doi: 10.1038/nrmicro2832

Follett, C. L., Dutkiewicz, S., Ribalet, F., Zakem, E., Caron, D., Armbrust, E. V., et al. (2022). Trophic interactions with heterotrophic bacteria limit the range of Prochlorococcus. Proc Natl Acad Sci U S A 119. doi: 10.1073/pnas.2110993118

Fuhrman, J. A. (2009). Microbial community structure and its functional implications. Nature 459, 193–199. doi: 10.1038/nature08058

Fuhrman, J. A., Cram, J. A., and Needham, D. M. (2015). Marine microbial community dynamics and their ecological interpretation. Nature Reviews Microbiology 13, 133–146. doi: 10.1038/nrmicro3417

Ghoul, M., and Mitri, S. (2016). The Ecology and Evolution of Microbial Competition. Trends in Microbiology 24, 833–845. doi: 10.1016/j.tim.2016.06.011

Giovannoni, S. J. (2017). SAR11 bacteria: The most abundant plankton in the oceans. Annu Rev Mar Sci 9, 231–255. doi: 10.1146/annurev-marine-010814-015934

Glenn, T. C., Pierson, T. W., Bayona-Vásquez, N. J., Kieran, T. J., Hoffberg, S. L., Thomas IV, J. C., et al. (2019). Adapterama II: universal amplicon sequencing on Illumina platforms (TaggiMatrix). PeerJ 7, e7786. doi: 10.7717/peerj.7786

González, A. M. M., Dalsgaard, B., and Olesen, J. M. (2010). Centrality measures and the importance of generalist species in pollination networks. Ecological Complexity 7, 36–43. doi: 10.1016/j.ecocom.2009.03.008

Guillou, L., Bachar, D., Audic, S., Bass, D., Berney, C., Bittner, L., et al. (2012). The Protist Ribosomal Reference database (PR2): a catalog of unicellular eukaryote Small Sub-Unit rRNA sequences with curated taxonomy. Nucleic Acids Research 41, D597–D604. doi: 10.1093/nar/gks1160

Guillou, L., Viprey, M., Chambouvet, A., Welsh, R. M., Kirkham, A. R., Massana, R., et al. (2008). Widespread occurrence and genetic diversity of marine parasitoids belonging to Syndiniales (Alveolata). Environ Microbiol 10, 3349–3365. doi: 10.1111/j.1462-2920.2008.01731.x

Hannigan, G. D., Duhaime, M. B., Koutra, D., and Schloss, P. D. (2018). Biogeography and environmental conditions shape bacteriophage-bacteria networks across the human microbiome. PLoS Comput Biol 14, e1006099. doi: 10.1371/journal.pcbi.1006099

Harrell Jr, F., and Dupont, C. (2019). Hmisc: Harrell Miscellaneous Version. Available at: https://CRAN.R-project.org/package=Hmisc

Harvell, C. D., Mitchell, C. E., Ward, J. R., Altizer, S., Dobson, A. P., Ostfeld, R. S., et al. (2002). Climate warming and disease risks for terrestrial and marine biota. Science 296, 2158–2162. doi: 10.1126/science.1063699

Hennon, G. M., Morris, J. J., Haley, S. T., Zinser, E. R., Durrant, A. R., Entwistle, E., et al. (2018). The impact of elevated CO(2) on Prochlorococcus and microbial interactions with “helper” bacterium Alteromonas. ISME J 12, 520–531. doi: 10.1038/ismej.2017.189

Hibbing, M. E., Fuqua, C., Parsek, M. R., and Peterson, S. B. (2010). Bacterial competition: surviving and thriving in the microbial jungle. Nat Rev Microbiol 8, 15–25. doi: 10.1038/nrmicro2259

Hoarfrost, A., Nayfach, S., Ladau, J., Yooseph, S., Arnosti, C., Dupont, C. L., et al. (2020). Global ecotypes in the ubiquitous marine clade SAR86. The ISME Journal 14, 178–188. doi: 10.1038/s41396-019-0516-7

Hofer, U. (2018). The majority is uncultured. Nature Reviews Microbiology 16, 716–717. doi: 10.1038/s41579-018-0097-x

Holen, D. (2014). Fishing for community and culture: the value of fisheries in rural Alaska. Polar Record 50, 403–413. doi: 10.1017/S0032247414000205

Holling, C. S. (1959). Some Characteristics of Simple Types of Predation and Parasitism. The Canadian Entomologist 91, 385–398. doi: 10.4039/Ent91385-7

Hu, H., Nigmatulina, K., and Eckhoff, P. (2013). The scaling of contact rates with population density for the infectious disease models. Mathematical Biosciences 244, 125–134. doi: 10.1016/j.mbs.2013.04.013

Hutchins, D. A., and Fu, F. (2017). Microorganisms and ocean global change. Nature Microbiology 2, 17058. doi: 10.1038/nmicrobiol.2017.58

Iida, Y., Hobley, L., Lambert, C., Fenton, A. K., Sockett, R. E., and Aizawa, S.-I. (2009). Roles of multiple flagellins in flagellar formation and flagellar growth post bdelloplast lysis in Bdellovibrio bacteriovorus. J Mol Biol 394, 1011–1021. doi: 10.1016/j.jmb.2009.10.003

Jeong, H., Tombor, B., Albert, R., Oltvai, Z. N., and Barabási, A.-L. (2000). The large-scale organization of metabolic networks. Nature 407, 651–654. doi: 10.1038/35036627

Jonsson, P. (1989). Photosynthetic assimilation of inorganic carbon in marine oligotrich ciliates (Ciliophora, Oligotrichina). Mar Microb Food Webs 2, 55–68.

Kassambara, A. (2020). ggplot2 Based Publication Ready Plots. Available at: https://rpkgs.datanovia.com/ggpubr/ (Accessed June 3, 2021).

Kodera, S. M., Das, P., Gilbert, J. A., and Lutz, H. L. (2022). Conceptual strategies for characterizing interactions in microbial communities. iScience 25, 103775. doi: 10.1016/j.isci.2022.103775

Koppelle, S., López-Escardó, D., Brussaard, C. P. D., Huisman, J., Philippart, C. J. M., Massana, R., et al. (2022). Mixotrophy in the bloom-forming genus Phaeocystis and other haptophytes. Harmful Algae 117, 102292. doi: 10.1016/j.hal.2022.102292

Ladd, C., Mordy, C., Kachel, N., and Stabeno, P. (2007). Northern Gulf of Alaska eddies and associated anomalies. Deep-Sea Research I 54, 487–509. doi: 10.1016/j.dsr.2007.01.006

Laloux, G. (2019). Shedding Light on the Cell Biology of the Predatory Bacterium Bdellovibrio bacteriovorus. Front Microbiol 10, 3136. doi: 10.3389/fmicb.2019.03136

Lambert, C., Evans, K. J., Till, R., Hobley, L., Capeness, M., Rendulic, S., et al. (2006). Characterizing the flagellar filament and the role of motility in bacterial prey-penetration by Bdellovibrio bacteriovorus. Mol Microbiol 60, 274–286. doi: 10.1111/j.1365-2958.2006.05081.x

Lambert, C., Fenton, A. K., Hobley, L., and Sockett, R. E. (2011). Predatory Bdellovibrio bacteria use gliding motility to scout for prey on surfaces. J Bacteriol 193, 3139–3141. doi: 10.1128/JB.00224-11

Li, L., Pujari, L., Wu, C., Huang, D., Wei, Y., Guo, C., et al. (2021). Assembly Processes and Co-occurrence Patterns of Abundant and Rare Bacterial Community in the Eastern Indian Ocean. Front Microbiol 12, 616956. doi: 10.3389/fmicb.2021.616956

Li, Y., Kan, J., Liu, F., Lian, K., Liang, Y., Shao, H., et al. (2024). Depth shapes microbiome assembly and network stability in the Mariana Trench. Microbiology Spectrum 12, e02110–23. doi: 10.1128/spectrum.02110-23

Lima-Mendez, G., Faust, K., Henry, N., Decelle, J., Colin, S., Carcillo, F., et al. (2015). Determinants of community structure in the global plankton interactome. Science 348. doi: 10.1126/science.1262073

Lin, Y.-C., Campbell, T., Chung, C.-C., Gong, G.-C., Chiang, K.-P., and Worden, A. Z. (2012). Distribution patterns and phylogeny of marine stramenopiles in the north pacific ocean. Appl Environ Microbiol 78, 3387–3399. doi: 10.1128/AEM.06952-11

Lok, C. (2015). Mining the microbial dark matter. Nature 522, 270–273. doi: 10.1038/522270a

Lucas, L., Koseff, J., Cloern, J., Monismith, S., and Thompson, J. (1999). Processes governing phytoplankton blooms in estuaries. I:The local production-loss balance. Marine Ecology Progress Series 187, 1–15. doi: 10.3354/meps187001

Massana, R., Unrein, F., Rodríguez-Martínez, R., Forn, I., Lefort, T., Pinhassi, J., et al. (2009). Grazing rates and functional diversity of uncultured heterotrophic flagellates. ISME J 3, 588–596. doi: 10.1038/ismej.2008.130

Matchado, M. S., Lauber, M., Reitmeier, S., Kacprowski, T., Baumbach, J., Haller, D., et al. (2021). Network analysis methods for studying microbial communities: A mini review. Comput Struct Biotechnol J 19, 2687–2698. doi: 10.1016/j.csbj.2021.05.001

McManus, G. B., Schoener, D., and Haberlandt, K. (2012). Chloroplast symbiosis in a marine ciliate: ecophysiology and the risks and rewards of hosting foreign organelles. Frontiers in Microbiology 3. doi: 10.3389/fmicb.2012.00321

Milici, M., Vital, M., Tomasch, J., Badewien, T. H., Giebel, H.-A., Plumeier, I., et al. (2017). Diversity and community composition of particle-associated and free-living bacteria in mesopelagic and bathypelagic Southern Ocean water masses: Evidence of dispersal limitation in the Bransfield Strait. Limnology and Oceanography 62, 1080–1095. doi: 10.1002/lno.10487

Mitulla, M., Dinasquet, J., Guillemette, R., Simon, M., Azam, F., and Wietz, M. (2016). Response of bacterial communities from California coastal waters to alginate particles and an alginolytic Alteromonas macleodii strain. Environmental Microbiology 18, 4369–4377. doi: 10.1111/1462-2920.13314

Montesinos-Navarro, A., Hiraldo, F., Tella, J. L., and Blanco, G. (2017). Network structure embracing mutualism–antagonism continuums increases community robustness. Nature Ecology & Evolution 1, 1661–1669. doi: 10.1038/s41559-017-0320-6

Mookherjee, A., and Jurkevitch, E. (2022). Interactions between Bdellovibrio and like organisms and bacteria in biofilms: beyond predator–prey dynamics. Environmental Microbiology 24, 998– 1011. doi: 10.1111/1462-2920.15844

Moreno, A. R., Anderson, C., Kudela, R. M., Sutula, M., Edwards, C., and Bianchi, D. (2022). Development, calibration, and evaluation of a model of Pseudo-nitzschia and domoic acid production for regional ocean modeling studies. Harmful Algae 118, 102296. doi: 10.1016/j.hal.2022.102296

Morris, J. J., Johnson, Z. I., Szul, M. J., Keller, M., and Zinser, E. R. (2011). Dependence of the Cyanobacterium Prochlorococcus on Hydrogen Peroxide Scavenging Microbes for Growth at the Ocean’s Surface. PLOS ONE 6, 1–13. doi: 10.1371/journal.pone.0016805

Morris, J. J., Kirkegaard, R., Szul, M. J., Johnson, Z. I., and Zinser, E. R. (2008). Facilitation of Robust Growth of *Prochlorococcus* Colonies and Dilute Liquid Cultures by “Helper” Heterotrophic Bacteria. Applied and Environmental Microbiology 74, 4530–4534. doi: 10.1128/AEM.02479-07

Morris, J. J., Lenski, R. E., and Zinser, E. R. (2012). The Black Queen Hypothesis: Evolution of Dependencies through Adaptive Gene Loss. mBio 3, e00036–12. doi: 10.1128/mBio.00036-12

Needham, D. M., Chow, C.-E. T., Cram, J. A., Sachdeva, R., Parada, A., and Fuhrman, J. A. (2013). Short-term observations of marine bacterial and viral communities: patterns, connections and resilience. The ISME Journal 7, 1274–1285. doi: 10.1038/ismej.2013.19

Negus, D., Moore, C., Baker, M., Raghunathan, D., Tyson, J., and Sockett, R. E. (2017). Predator Versus Pathogen: How Does Predatory Bdellovibrio bacteriovorus Interface with the Challenges of Killing Gram-Negative Pathogens in a Host Setting? Annual Review of Microbiology 71, 441–457. doi: 10.1146/annurev-micro-090816-093618

Newman, M. E. J. (2003). A measure of betweenness centrality based on random walks.

O’Hara, M. (2023). Distribution and Mixotrophy of Cryptophyte Phytoplankton in the Northern Gulf of Alaska. Western Washington University. Available at: https://books.google.com/books?id=EDbLzwEACAAJ

Okkonen, S., Jacobs, G., Metzger, E., Hurlburt, H., and Shriver, J. (2001). Mesoscale variability in the boundary currents of the Alaska Gyre. Continental Shelf Research 21, 1219–1236. doi: 10.1016/S0278-4343(00)00085-6

Ortega, E. L. S., Reister, I., Danielson, S. L., and Aguilar-Islas, A. M. (2025). Surface macro- and micro-nutrients within the Copper River plume region respond to along-shore winds. Marine Chemistry 270, 104508. doi: 10.1016/j.marchem.2025.104508

Parada, A. E., Needham, D. M., and Fuhrman, J. A. (2016). Every base matters: assessing small subunit rRNA primers for marine microbiomes with mock communities, time series and global field samples. Environmental Microbiology 18, 1403–1414. doi: 10.1111/1462-2920.13023

Poisot, T., Lepennetier, G., Martinez, E., Ramsayer, J., and Hochberg, M. E. (2011). Resource availability affects the structure of a natural bacteria–bacteriophage community. Biology Letters 7, 201–204. doi: 10.1098/rsbl.2010.0774

Qian, J. J., and Akçay, E. (2020). The balance of interaction types determines the assembly and stability of ecological communities. Nature Ecology & Evolution 4, 356–365. doi: 10.1038/s41559-020-1121-x

Quast, C., Pruesse, E., Yilmaz, P., Gerken, J., Schweer, T., Yarza, P., et al. (2013). The SILVA ribosomal RNA gene database project: improved data processing and web-based tools. Nucleic Acids Res 41, D590–D596. doi: 10.1093/nar/gks1219

Rial, P., Garrido, J. L., Jaén, D., and Rodríguez, F. (2012). Pigment composition in three Dinophysis species (Dinophyceae) and the associated cultures of Mesodinium rubrum and Teleaulax amphioxeia. Journal of Plankton Research 35, 433–437. doi: 10.1093/plankt/fbs099

Roda-Garcia, J. J., Haro-Moreno, J. M., Huschet, L. A., Rodriguez-Valera, F., and López-Pérez, M. (2021). Phylogenomics of SAR116 Clade Reveals Two Subclades with Different Evolutionary Trajectories and an Important Role in the Ocean Sulfur Cycle. mSystems 6, e0094421. doi: 10.1128/mSystems.00944-21

Rusterholz, P. M., Hansen, P. J., and Daugbjerg, N. (2017). Evolutionary transition towards permanent chloroplasts? - Division of kleptochloroplasts in starved cells of two species of Dinophysis (Dinophyceae). PLoS ONE 12, e0177512. doi: 10.1371/journal.pone.0177512

Sakka, A., Legendre, L., Gosselin, M., and Delesalle, B. (2000). Structure of the oligotrophic planktonic food web under low grazing of heterotrophic bacteria: Takapoto Atoll, French Polynesia. Mar Ecol Prog Ser 197, 1–17.

Sarmento, H., Romera-Castillo, C., Lindh, M., Pinhassi, J., Sala, M. M., Gasol, J. M., et al. (2013). Phytoplankton species-specific release of dissolved free amino acids and their selective consumption by bacteria. Limnology and Oceanography 58, 1123–1135. doi: 10.4319/lo.2013.58.3.1123

Scheffer, M., Carpenter, S. R., Lenton, T. M., Bascompte, J., Brock, W., Dakos, V., et al. (2012). Anticipating Critical Transitions. Science 338, 344–348. doi: 10.1126/science.1225244

Schlitzer, R. (2022). Ocean Data View. Available at: https://odv.awi.de/

Screen, J. A., and Simmonds, I. (2010). The central role of diminishing sea ice in recent Arctic temperature amplification. Nature 464, 1334–1337. doi: 10.1038/nature09051

Serreze, M. C., Barrett, A. P., Stroeve, J. C., Kindig, D. N., and Holland, M. M. (2009). The emergence of surface-based Arctic amplification. The Cryosphere 3, 11–19. doi: 10.5194/tc-3-11-2009

Shannon, P., Markiel, A., Ozier, O., Baliga, N. S., Wang, J. T., Ramage, D., et al. (2003). Cytoscape: A Software Environment for Integrated Models of Biomolecular Interaction Networks. Genome Res. 13, 2498–2504. doi: 10.1101/gr.1239303

Šimek, K., Mukherjee, I., Szöke-Nagy, T., Haber, M., Salcher, M. M., and Ghai, R. (2023). Cryptic and ubiquitous aplastidic cryptophytes are key freshwater flagellated bacterivores. The ISME Journal 17, 84–94. doi: 10.1038/s41396-022-01326-4

Sockett, R. E. (2009). Predatory Lifestyle of Bdellovibrio bacteriovorus. Annu. Rev. Microbiol. 63, 523–539. doi: 10.1146/annurev.micro.091208.073346

Stabeno, P. J., Bell, S., Cheng, W., Danielson, S., Kachel, N. B., and Mordy, C. W. (2016). Long-term observations of Alaska Coastal Current in the northern Gulf of Alaska. Deep Sea Research Part II: Topical Studies in Oceanography 132, 24–40. doi: 10.1016/j.dsr2.2015.12.016

Stabeno, P. J., Bond, N. A., Hermann, A. J., Kachel, N. B., Mordy, C. W., and Overland, J. E. (2004). Meteorology and oceanography of the Northern Gulf of Alaska. Continental Shelf Research 24, 859–897. doi: 10.1016/j.csr.2004.02.007

Stoecker, D. K., Hansen, P. J., Caron, D. A., and Mitra, A. (2017). Mixotrophy in the Marine Plankton. Annual Review of Marine Science 9, 311–335. doi: 10.1146/annurev-marine-010816-060617

Stoecker, D. K., and Lavrentyev, P. J. (2018). Mixotrophic Plankton in the Polar Seas: A Pan-Arctic Review. Frontiers in Marine Science 5. doi: 10.3389/fmars.2018.00292

Stoecker, D. K., and Silver, M. W. (1990). Replacement and aging of chloroplasts inStrombidium capitatum (Ciliophora: Oligotrichida). Marine Biology 107, 491–502. doi: 10.1007/BF01313434

Stoecker, D. K., Silver, M. W., Michaels, A. E., and Davis, L. H. (1988). Obligate mixotrophy inLaboea strobila, a ciliate which retains chloroplasts. Marine Biology 99, 415–423. doi: 10.1007/BF02112135

Strom, S. (2002). Novel interactions between phytoplankton and microzooplankton: their influence on the coupling between growth and grazing rates in the sea. Hydrobiologia 480, 41–54. doi: 10.1023/A:1021224832646

Strom, S. (2023). Recent Marine Heatwaves Affect Marine Ecosystems from Plankton to Seabirds in the Northern Gulf of Alaska. Oceanog. doi: 10.5670/oceanog.2023.s1.9

Strom, S., Bright, K., and Fredrickson, K. (2024). Widespread ciliate and dinoflagellate mixotrophy may contribute to ecosystem resilience in a subarctic sea: the northern Gulf of Alaska. Aquat Microb Ecol 90, 1–21.

Strom, S. L., Macri, E. L., and Fredrickson, K. A. (2010). Light limitation of summer primary production in the coastal Gulf of Alaska: physiological and environmental causes. Marine Ecology Progress Series 402, 45–57.

Strom, S., Olson, M., Macri, E., and Mord, C. (2006). Cross-shelf gradients in phytoplankton community structure, nutrient utilization, and growth rate in the coastal Gulf of Alaska. Mar. Ecol. Prog. Ser. 328, 75–92. doi: 10.3354/meps328075

Szymkowiak, M. (2020). Adaptations and well-being: Gulf of Alaska fishing families in a changing landscape. Ocean & Coastal Management 197, 105321. doi: 10.1016/j.ocecoaman.2020.105321

Thébault, E., and Fontaine, C. (2010). Stability of Ecological Communities and the Architecture of Mutualistic and Trophic Networks. Science 329, 853–856. doi: 10.1126/science.1188321

Thingstad, T. F. (2000). Elements of a theory for the mechanisms controlling abundance, diversity, and biogeochemical role of lytic bacterial viruses in aquatic systems. Limnology and Oceanography 45, 1320–1328. doi: 10.4319/lo.2000.45.6.1320

Thingstad, T. F., and Lignell, R. (1997). Theoretical models for the control of bacterial growth rate, abundance, diversity and carbon demand. Aquatic Microbial Ecology - AQUAT MICROB ECOL 13, 19–27. doi: 10.3354/ame013019

Thingstad, T. F., Våge, S., Storesund, J. E., Sandaa, R.-A., and Giske, J. (2014). A theoretical analysis of how strain-specific viruses can control microbial species diversity. Proceedings of the National Academy of Sciences 111, 7813–7818. doi: 10.1073/pnas.1400909111

Tsuchiya, M., Miyawaki, S., Oguri, K., Toyofuku, T., Tame, A., Uematsu, K., et al. (2020). Acquisition, Maintenance, and Ecological Roles of Kleptoplasts in Planoglabratella opercularis (Foraminifera, Rhizaria). Frontiers in Marine Science 7. doi: 10.3389/fmars.2020.00585

Tucker, S. J., Freel, K. C., Monaghan, E. A., Sullivan, C. E. S., Ramfelt, O., Rii, Y. M., et al. (2021). Spatial and temporal dynamics of SAR11 marine bacteria across a nearshore to offshore transect in the tropical Pacific Ocean. PeerJ 9, e12274. doi: 10.7717/peerj.12274

Wang, F., Li, M., Huang, L., and Zhang, X.-H. (2021). Cultivation of uncultured marine microorganisms. Marine Life Science & Technology 3, 117–120. doi: 10.1007/s42995-021-00093-z

Wang, P., Li, S.-P., Yang, X., Si, X., Li, W.-J., Shu, W., et al. (2023). Spatial scaling of soil microbial co-occurrence networks in a fragmented landscape. mLife 2, 209–215. doi: 10.1002/mlf2.12073

Wang, Z., Juarez, D. L., Pan, J.-F., Blinebry, S. K., Gronniger, J., Clark, J. S., et al. (2019). Microbial communities across nearshore to offshore coastal transects are primarily shaped by distance and temperature. Environmental Microbiology 21, 3862–3872. doi: 10.1111/1462-2920.14734

Weiss, S., Van Treuren, W., Lozupone, C., Faust, K., Friedman, J., Deng, Y., et al. (2016). Correlation detection strategies in microbial data sets vary widely in sensitivity and precision. ISME J 10, 1669–1681. doi: 10.1038/ismej.2015.235

Weitz, J. S., and Dushoff, J. (2008). Alternative stable states in host–phage dynamics. Theoretical Ecology 1, 13–19. doi: 10.1007/s12080-007-0001-1

Widder, S., Besemer, K., Singer, G. A., Ceola, S., Bertuzzo, E., Quince, C., et al. (2014). Fluvial network organization imprints on microbial co-occurrence networks. Proc Natl Acad Sci U S A 111, 12799–12804. doi: 10.1073/pnas.1411723111

Winter, C., Bouvier, T., Weinbauer, M. G., and Thingstad, T. F. (2010). Trade-offs between competition and defense specialists among unicellular planktonic organisms: the “killing the winner” hypothesis revisited. Microbiol Mol Biol Rev 74, 42–57. doi: 10.1128/MMBR.00034-09

Worden, A. Z., Follows, M. J., Giovannoni, S. J., Wilken, S., Zimmerman, A. E., and Keeling, P. J. (2015). Environmental science. Rethinking the marine carbon cycle: factoring in the multifarious lifestyles of microbes. Science 347, 1257594. doi: 10.1126/science.1257594

Yan, C., Owen, J. S., Seo, E.-Y., Jung, D., and He, S. (2023). Microbial Interaction is Among the Key Factors for Isolation of Previous Uncultured Microbes. J Microbiol 61, 655–662. doi: 10.1007/s12275-023-00063-3

Yang, M., Zhao, L., Yu, X., Shu, W., Cao, F., Liu, Q., et al. (2024). Microbial community structure and co-occurrence network stability in seawater and microplastic biofilms under prometryn pollution in marine ecosystems. Mar Pollut Bull 199, 115960. doi: 10.1016/j.marpolbul.2023.115960

Yang, Y., Shi, Y., Fang, J., Chu, H., and Adams, J. M. (2022). Soil Microbial Network Complexity Varies With pH as a Continuum, Not a Threshold, Across the North China Plain. Frontiers in Microbiology 13. doi: 10.3389/fmicb.2022.895687

Zelezniak, A., Andrejev, S., Ponomarova, O., Mende, D. R., Bork, P., and Patil, K. R. (2015). Metabolic dependencies drive species co-occurrence in diverse microbial communities. Proceedings of the National Academy of Sciences 112, 6449–6454. doi: 10.1073/pnas.1421834112

Zhang, S., Zheng, T., Lundholm, N., Huang, X., Jiang, X., Li, A., et al. (2021). Chemical and morphological defenses of Pseudo-nitzschia multiseries in response to zooplankton grazing. Harmful Algae 104, 102033. doi: 10.1016/j.hal.2021.102033

