## Supplementary figures and images for "Identifying potential keystone microbes from co-occurrence networks in the Gulf of Alaska"

### Supplemental Figure 1

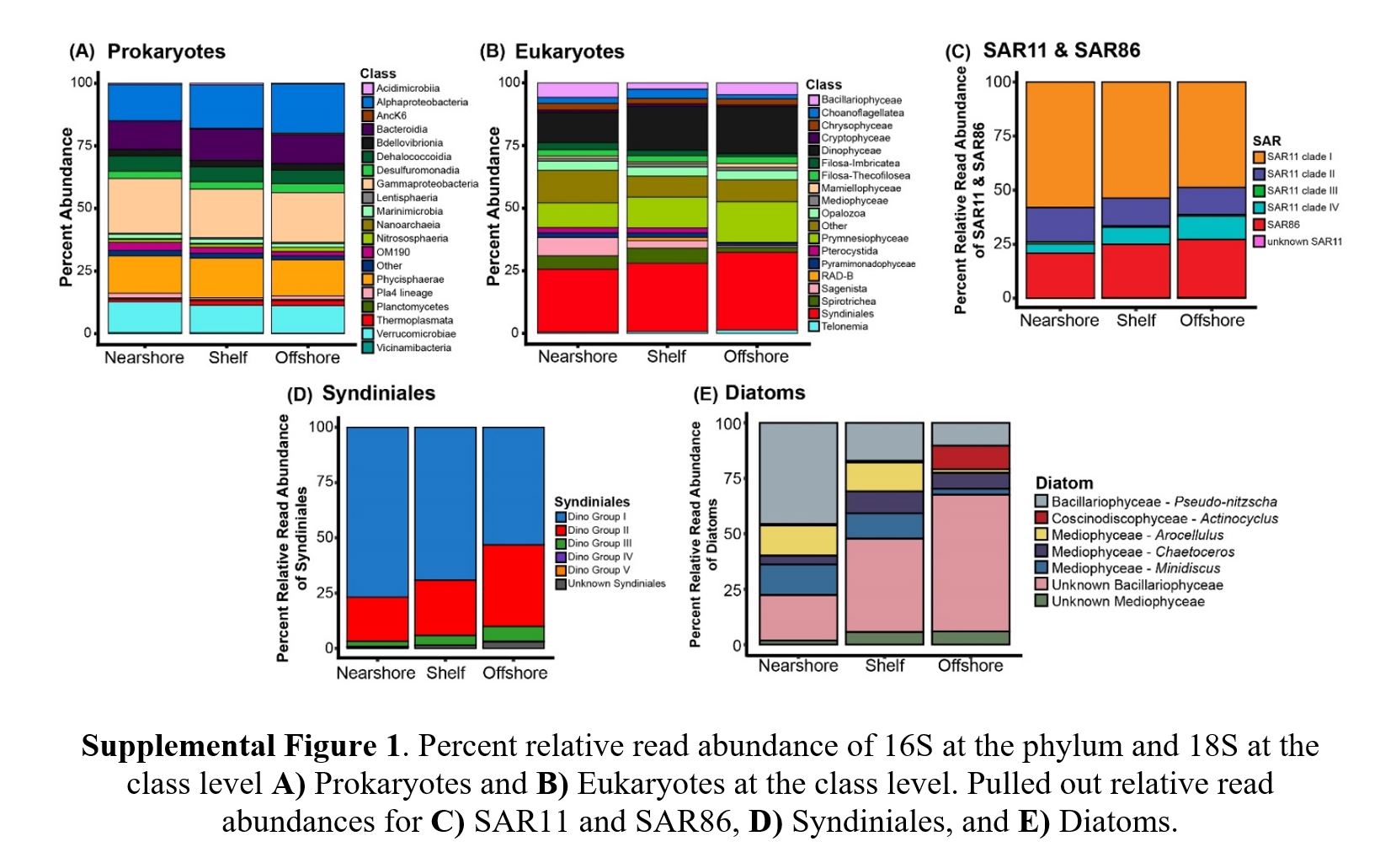

### Supplemental Figure 2

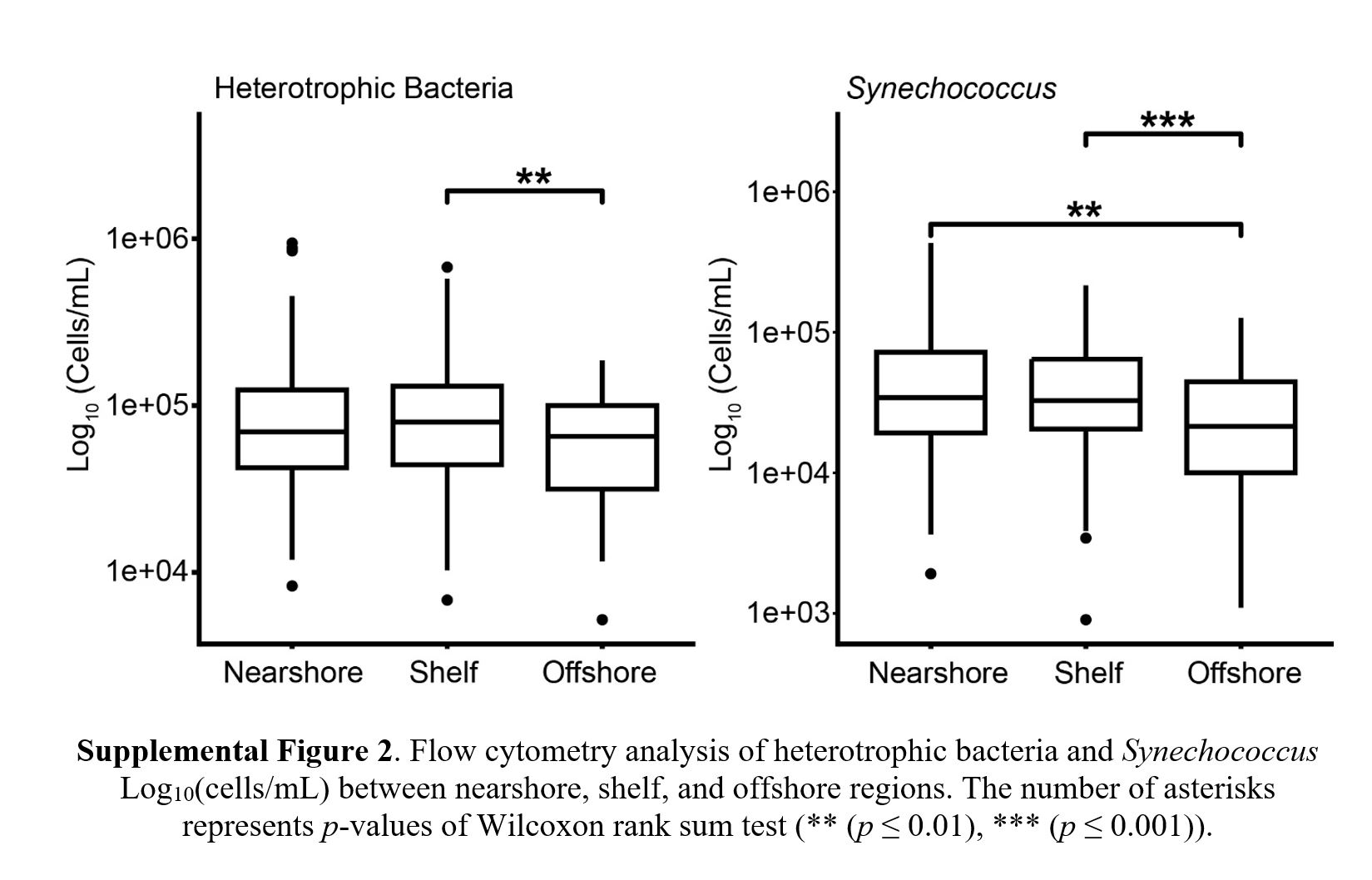
