## Supplemental Table 1 for "Identifying potential keystone microbes from co-occurrence networks in the Gulf of Alaska"

**Supplemental Table 1**. Sample counts and locations

| Region | 16S Sample Count | 18S Sample Count |
| --- | --- | --- |
| Nearshore | 43 | 33 |
| Shelf | 74 | 60 |
| Offshore | 58 | 52 |
|  | 175 | 145 |

|  | 2018 | 2019 | 2020 | 2021 |
| --- | --- | --- | --- | --- |
| Seward Line | 9 | 30 | 24 | 29 |
| Middleton Island Line | 8 | 19 | 15 | 21 |
| Kodiak Island Line | 9 | 19 | 0 | 17 |
