## Supplemental Table 2 for "Identifying potential keystone microbes from co-occurrence networks in the Gulf of Alaska"

**Supplemental Table 2**. Network statistic definitions.

| Term | Definition |
| --- | --- |
| Node-specific | |
| Average shortest path | The average length of the shortest path between the node and all other nodes. This measures the overall efficiency of the network in terms of how quickly information could be transmitted between nodes. |
| Closeness Centrality | The reciprocal of the average shortest path length. A measure of how close a node is to all other nodes in the network. Nodes with higher closeness centrality are generally more "central" and can spread information more efficiently. |
| Degree Centrality | The number of edges linked to each node. It is an indicator of a node's immediate influence or activity within the network. Higher degree centrality means more connections. |
| Edge | A significant Spearman correlation (ρ > 0.8) between two nodes. |
| Node | An operational taxonomic unit with significant correlations. |
| Global Network | |
| Average number of neighbors | The average number of direct edges per node in the network. It gives an idea of how connected each node is on average within the network. |
| Centralization | A measure of how centralized the network is, based on the centrality of its nodes. A highly centralized network has one or a few nodes that dominate the network, while a decentralized network has nodes with more equal centrality. |
| Clustering coefficient | A measure of the degree to which nodes in the network tend to cluster together. It reflects the likelihood that the neighbors of a node are also connected to each other. High clustering indicates microbial species likely form tightly-knit groups. |
| Components | Subnetworks within the larger network that are connected internally but are not connected to other parts of the network. A network might have multiple components if it is not fully connected. |
| Density | The ratio of edges in the network to the total number of possible edges. It measures how densely connected the network is. A high density indicates many interactions between nodes relative to the total possible interactions. |
| Diameter | The longest shortest path between any two nodes in the network. It represents the maximum distance between the farthest nodes and gives an indication of the "spread" of the network. |
| Fragmentation | The ratio of disconnected subgraphs (CL) to the overall number of nodes (N) in each network. A measure of the extend to which the network is broken up into smaller disconnected parts or components. High fragmentation means that the network has many isolated nodes or clusters. |
| Heterogeneity | The degree of variation in the network, particularly in terms of node degree centrality. A heterogeneous network has nodes with very different numbers of connections, whereas a homogeneous network has nodes with more similar degrees |
| Path Length | The length of the shortest path between two nodes. It is used to quantify the distance between nodes in terms of the number of edges traversed. |
| Radius | The smallest distance across all nodes and indicates the "core" of the network. |
